# Learning, Visualizing and Exploring 16S rRNA Structure Using an Attention-based Deep Neural Network

**DOI:** 10.1101/2020.10.12.336271

**Authors:** Zhengqiao Zhao, Stephen Woloszynek, Felix Agbavor, Joshua Chang Mell, Bahrad A. Sokhansanj, Gail Rosen

## Abstract

Recurrent neural networks (RNNs) with memory (e.g. LSTMs) and attention mechanisms are widely used in natural language processing because they can capture short and long term sequential information for diverse tasks. We propose an integrated deep learning model for microbial DNA sequence data, which exploits convolutional networks, recurrent neural networks, and attention mechanisms to perform sample-associated attribute prediction—*phenotype prediction*—and extract interesting features, such as informative taxa and predictive *k*-mer context. In this paper, we develop this novel deep learning approach and evaluate its application to amplicon sequences. We focus on typically short DNA reads of 16s ribosomal RNA (rRNA) marker genes, which identify the heterogeneity of a microbial community sample. Our deep learning approach enables sample-level attribute and taxonomic prediction, with the aim of aiding biological research and supporting medical diagnosis. We demonstrate that our implementation of a novel attention-based deep network architecture, `Read2Pheno`, achieves read-level phenotypic prediction and, in turn, that aggregating read-level information can robustly predict microbial community properties, host phenotype, and taxonomic classification, with performance comparable to conventional approaches. Most importantly, as a further result of the training process, the network architecture will encode sequences (reads) into dense, meaningful representations: learned embedded vectors output on the intermediate layer of the network model, which can provide biological insight when visualized. Finally, we demonstrate that a model with an attention layer can automatically identify informative regions in sequences/reads which are particularly informative for classification tasks. An implementation of the attention-based deep learning network is available at https://github.com/EESI/sequence_attention.

## Introduction

Advances in DNA sequencing are rapidly producing complex microbiome datasets in fields ranging from human health to environmental studies [1]. Large-scale microbial projects provide rich information, enabling the prediction of sample-level traits (i.e., phenotypes), aiding in biological discovery, and supporting medical diagnosis. A typical microbiome study may contain hundreds to thousands of samples. In turn, each sample contains thousands of reads depending on the sequence depth. These reads are fragments of DNA/RNA material extracted from microbes residing in the environment where the sample was collected. Hence, by way of example, an environmental sample can be sequenced via 16S ribosomal RNA amplicon technology, to provide a comprehensive taxonomic survey of an environment’s or subject’s microbial community [2, 3]. Recently, the decrease in cost of next-generation high throughput sequencing technology has further allowed the use of metagenomic approaches, such as shotgot sequencing to generate a dataset that reflects both taxonomy and gene sequences [3–5].

A major focus of microbiome research has been, and continues to be, the use of 16s rRNA amplicon sequencing surveys to determine “Who is there?” in a host or environmental sample. The answer to “Who is there?” may, in turn, be used to predict host phenotype for clinical diagnoses or infer taxa-phenotype association for basic biology research [6–10]. In the context of our work, we define “phenoytpe” as an overall trait at the environmental level or habitat that the microbiome sample is isolated from [11, 12], thereby incorporating the emergent function of the microbiome (a.k.a. microbiome phenotypes) [13–18]. For example, the expansive definition of “phenotype” in the microbiome context can include the preference of a certain microbial community for a particular environmental niche or body site [19]. Thus, the microbiome may be shaped by the environment.

### 16S rRNA marker gene-based phenotype prediction

Analyzing “Who is there?” through 16s rRNA amplicon sequencing is relatively affordable and easy to implement in the field—but *phenotype prediction* from rRNA sequence is a major challenge. Ribosomal sequence does not itself contain functional information, unlike, e.g., more costly and complex metagenomic shotgun sequencing data [9, 20]). Building machine learning phenotype classifiers usually starts with constructing a microbial abundance table, such as an Operational Taxonomic Unit (OTU) table, an Amplicon Sequence Variant (ASV) table, or a *k*-mer frequency table (i.e., table of the frequencies of k-length nucleotide strings within the collection of reads in a sample) [8, 9]. Researchers then train a classifier to distinguish phenotypes by learning from the taxon abundance of sequenced samples in a training data set. For example, a classifier may be constructed to identify a sample as being from the gut from a patient diagnosed with a disease. In this example, if a certain combination of some taxa in a novel sample are more abundant than a threshold previously determined based on a training data set of gut samples, the novel sample will be identified as disease-positive.

By analyzing the OTU/ASV abundance table, therefore, researchers can discover underlying associations between certain taxa or groups of taxa and phenotype. For example, in Gevers *et al*. [7], samples were collected from a) patients with Crohn’s disease and b) control groups. Gevers *et al*. discovered some bacterial taxa which were solely abundant in disease groups, along with some taxa which were eliminated by infection of the disease. These findings are helpful in disease diagnoses and treatment. A systematic survey of 18 classification methods and 5 feature selection methods were assessed to classify phenotypes [21]. To do so, the authors transformed the 16S rRNA sequences to OTU tables, which in turn served as the input to the algorithms under evaluation. The authors showed that feature selection can improve phenotype prediction performance for many classification algorithms, and that Random Forests with optimized parameters are nominally the best performing classifier. Another phenotype prediction method that has been proposed and evaluated is the RoDEO normalization (Robust Differential Expression Operator) based classifier [8]. In [8], the authors compared various normalization methods for OTU data and showed that RoDEO processed count data with linear kernel support vector machines produced the best performance on multiple experimental datasets. The authors further showed that using a small subset of OTUs sometimes gave better accuracy than using all OTUs.

The construction of OTU/ASV tables, however, often involves denoising, sequence alignment, and taxonomic classification, and thus can lead to information loss from the true information contained in the raw nucleotide reads. By grouping sequences to limited taxonomic labels, it becomes difficult to quantify the genotype-to-phenotype relationship. Of particular concern is the omission of nucleotide structural information from OTU mapping, where the 97% identity threshold conventionally used for OTU mapping smooths over valuable nucleotide variation. This is better addressed through the more exact ASV identification—but rarely is the nucleotide level information examined past the mapping step. Alternatively, a *k*-mer representation of amplicon sequences has been proposed to predict phenotype, which is shown can outperform traditional OTU representation [9]. Since a *k*-mer-based method is alignment free and reference free, it would cost less computationally than OTU-based methods. Because *k*-mer representations cut reads into smaller pieces, methods based on *k*-mers will lose sequential information. As such, *k*-mer analysis is subject to the length of the *k*-mers and does not preserve the nucleotide context/sequential-order. Some local nucleotide variation may be able to be identified; however, the long-range nucleotide sequential information is completely lost. In sum, currently available methods are unable to easily and robustly connect nucleotide changes on the read level back to the phenotype prediction and thereby reveal which nucleotide features are specifically relevant to the classification.

### Deep neural networks and their application in bioinformatics

Recent advances in supervised deep learning are further able to leverage a huge volume of different kinds of data. Convolutional neural networks (CNNs), which may be interpreted by saliency maps [22], have been vital to image recognition. Model interpretability, in general, has been a research direction of particular interest in the deep learning field [23–25]. Successes in these other areas have inspired applications of deep learning to bioinformatics as well [26]. In addition, deep learning approaches can learn hierarchical representations of metagenomic data that standard classification methods do not allow [27]. Both CNNs and RNNs have been applied to areas such as transcription factor binding site classification [28, 29], SNP calling [30, 31], microbial taxonomic classification [32] and DNA sequence function prediction and gene inference [33, 34]. The authors explore deep learning approaches for predicting environments and host phenotype using *k*-mer-based representation of shallow subsamples in [9]. Lo *et al*. proposed deep learning approaches to learn microbial count data (e.g., OTU table) for host phenotypes prediction [35]. The microbial count data can be formatted into an “image” format to be processed by a CNN model [36]. The CNN model has also been used to learn phylogenetic structure of a metagenomic sample to predict the host phenotype [37]. In this work, a 2D matrix is used to represent the phylogenetic tree of microbial taxa (with relative abundance) in a sample, and a CNN model is designed to learn from such data. Woloszynek *et al*. proposed an unsupervised method to embed 16S rRNA sequences to meaningful numerical vectors to facilitate the down-stream analysis and visualization [38]. Many models rely on extracting “features” (for instance, taxonomic composition or functional profiles) from the sequence data [39].

### Exploring connections between genetic features and biological classes

In addition to making predictions, machine learning models can reveal knowledge about domain relationships contained in data, often referred to as interpretations [40]. In the context of sequence classification tasks, i.e., microbial survey data based phenotype prediction, once a predictive model is built, the researchers can further identify sequence features relevant to classifications, i.e., occurring taxa and gnomic content related to a certain disease. There are substantial research attempts to identify label associated genetic content. A complementary approach is *supervised* computational method, as a means of associating genetic content with known labels, i.e., taxa. “Oligotyping” has been proposed as a way to identify subtypes of 16S rRNA sequence variation, based on distinguishing sequence variants by subsets of several nucleotides within the sequence, i.e., oligomers. Specifically, *Oligotyping* is a supervised computational method that identifies those nucleotide positions that represent information-rich variation [41]. *Oligotyping* requires information about the taxonomic classification of the sequence via OTU clustering or supervised methods. Then, the method is applied to taxonomical/OTU groups of interest. *Oligotyping* can be an efficient way to identify meaningful subpopulations of a single species and informative nucleotides. However, preprocessing steps are still needed (e.g., OTU clustering or multiple sequence alignment) to find closely related sequences.

Another proposed method, “PhenotypeSeeker” [42], is a statistics-based framework to find genotype-phenotype associations. Predictive *k*-mers are identified by a regression model, and the statistical test further quantifies their relative importance. However, the authors only built species-specific models trained for a closely related group of bacterial isolates and their associated phenotypes. Visualization methods are developed for DNA/RNA binding sites prediction models as mentioned in Section Deep neural networks and their application in bioinformatics [28, 29, 43, 44] to reveal predictive genomic content. Alipanahi *et al*. propose to interpret the model and visualize informative single nucleotide polymorphisms (SNPs) by manually altering nucleotides in the input reads and comparing the resulting new prediction with the original prediction of the unaltered input [43]. In `Deep Motif`, the authors use Saliency Maps [22, 25] to interpret the model and visualize informative genomic content [28].

### Better interpretability: Attention mechanisms

Attention mechanisms have become more widely applied in the natural language processing (NLP) and image recognition fields to improve the interpretability of deep learning models [45–48]. For example, it has been shown that an attention-based Bi-LSTM (Bi-directional long short term memory) RNN model can successfully capture the most important semantic information in a sentence and outperform most existing competing approaches [47]. A hierarchical attention network can also improve document level classification [46] by selecting qualitatively informative words and sentences. Informative content may be visualized by looking at the output of the attention layers of the network model. The use of deep learning with attention mechanisms has also been suggested for the field of bioinformatics. Deming *et al*. [29] proposed a method for simultaneously learning general genomic patterns and identifies the sequence motifs that contribute most to predicting functional genomic outcomes. While they found a marked gain in performance over previous architectures, their model was not used for phenotype prediction.

In this paper, we exploit CNNs, RNNs, and attention mechanisms for phenotype/taxonomic prediction and propose a `Read2Pheno` classifier to predict phenotype from 16S rRNA reads and, thereby, explore and visualize informative nucleotide structure and taxa. The sample-to-phenotype prediction can then be inferred by a sample-level predictor which aggregates the abstraction of all reads from the `Read2Pheno` model1. We show that the model trained with read level information can achieve similar sample-to-phenotype predictions compared with conventional methods. We further provide a visualization of the embedded vectors, which is a representation of the information that the network is learning. We use attention weights to identify and visualize the nucleotides associated with phenotype and/or taxonomy, and compare the highlighted informative regions against a base-line entropy method and *Oligotyping* [41].

We show the efficacy of our model with the American Gut microbiome dataset [49] (http://americangut.org/), Gevers *et al*.’s Crohn’s disease dataset [7] and SILVA 16S rRNA dataset [50, 51] and explore interesting visualizations and features generated by the model. The experimental results show that the performance of our model is comparable to current methods and our models can provide further interpretation and visualization.

## Methods

Our proposed model consists of two parts: the `Read2Pheno` read-level classifier and the sample-level predictor. We first train a read-level classifier using an attention-based deep neural network to predict DNA/RNA reads to the sample level labels the reads associated with. For example, if the samples are labeled with collected body sites, the model will be trained to learn the original body site that the reads were collected from. Then, a sample-level prediction can be made by three different ways: 1) tally a majority vote of all the read prediction scores in the sample of interest to obtain a final prediction; 2) use the output of the intermediate layer to obtain a read embedding (see Figure 1 for details) and average read embeddings from a sample to gain an overall sample-level embedding that a classifier can train on to predict a sample-level label; 3) apply clustering on read embeddings of training data and assign reads per sample to those clusters to form a “pseudo” OTU table [38]. Then a classifier can be trained for phenotype prediction. In sum, our `Read2Pheno` read-level classifier can capture read level model and provide biological insights by read embedding ordination and attention weights visualization and the sample-level predictor can aggregate information learned in read-level and make sample-level classifications to validate our overall framework.

**Figure 1.**
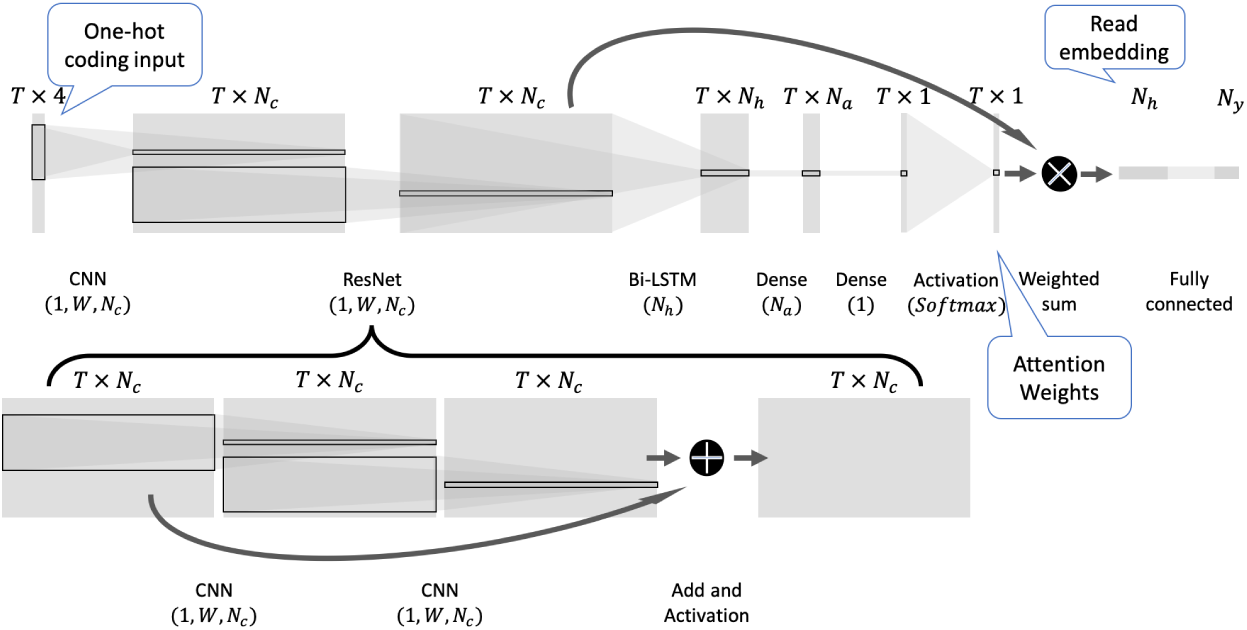
`Read2Pheno` classifier architecture: The input is a one-hot coded 16S rRNA sequence with length *T*. The input is fed to a few 1-dimentional convolutional blocks with window size of *W* and the number of output channels of *N*_*c*_. The resultant output is a *N*_*c*_ × *T* dimensional matrix which is then fed to a Bidirectional LSTM layer with the number of hidden nodes of *N*_*h*_. *N*_*a*_ is the number of hidden nodes used to compute attention weights and *N*_*y*_ is the total number of phenotypes (number of classes) to be predicted. There are two informative intermediate layer outputs (attention weights and read embedding vectors) which are labeled by blue tags. They are used in the analysis described in this paper.

### Read2Pheno Classifier

The `Read2Pheno` classifier is a hybrid convolutional and recurrent deep neural network with attention. Figure 1 shows a diagram of the classifier. Sequencing data are one-hot coded according to the map shown in Appendix One-hot coding map. Then the array representation of a read is fed into several initial layers of convolutional blocks (inspired by the scheme in [29]). The result is a embedding of the read, a *N*_*c*_ × *T* dimensional matrix, by learning local *k*-mer patterns, where *N*_*c*_ is the number of output channels in convolutional blocks and *T* is the length of input DNA reads. A Bi-directional Long Short Term Memory (Bi-LSTM) model is then applied to the data to learn the longitude dependency of nucleotides. The returned sequence is then processed and normalized by an attention layer to get an attention vector using the soft attention mechanism, as described in [47, 52]. The output of Bi-LSTM layer in our model is a *N*_*h*_ × *T* dimensional matrix where *N*_*h*_ is the number of hidden nodes in Bi-LSTM layer and *T* is the length of input DNA reads. Each base position (time-step) in the input corresponds to a *N*_*h*_ dimensional vector (hidden states at this position). The dense attention layer applies to the hidden states of every base position (time-step). The dense layer thereby learns the importance of hidden states at each position and return a small value if the hidden states of this position do not make an important contribution to the model’s final prediction, and, conversely, a large value if the model relies on the hidden states at this position in making the final prediction. The output of the dense layer is a vector of length *T*. Then, the output is normalized by a `softmax` function to produce the attention vector [52]. The output of this layer naturally indicates the regions in the sequence that the model pays attention to. While the attention weights are not learned from specific nucleotides but from high level features from 9-mers and their sequential information, as shown in Figure 1, the attention interpretation may be considered to be an approximation of the informative nucleotides of the 16S rRNA gene. The final embedding of the read is a weighted sum of all the embeddings across the sequence, where the weights are the elements of the attention vector. The goal of this layer is to suppress the regions that are less relevant to prediction and focus on informative regions. Finally, a dense layer with `softmax` activation function is applied to the read embedding vector to classify it into one of *N*_*y*_ labels. The hyperparameter selection process is described in Section Model selection on American Gut dataset.

### Sample-level predictor

In this paper, we propose three different ways to perform sample-level prediction. The most intuitive way is performing a majority vote. The sample-level predictor counts all the votes, i.e., the resulting `Read2Pheno` classifications, from all the reads in a query sample and labeling the sample with the overall majority vote. The majority vote is a baseline method intended to illustrate that the `Read2Pheno` model is learning the sample-associated phenotypic labels for each read. We compare the majority vote baseline to proposed embedding-based approaches further described below.

The intermediate layer of our model provides a concise numerical representation of the input reads, which we can exploit in sample-level prediction. We propose to use two embedding based approaches: sample-level embedding method and “Pseudo OTU” method [38]. The sample-level embedding method forms a sample-level vector representation by averaging all read-level embeddings in a query sample. Then, a classifier, such as Random Forest, can be trained to learn the sample-phenotype association. For the “pseudo OTU” method as described by Woloszynek *et al*. [38], first read-level embedding vectors are clustered via an unsupervised algorithm such as k-means to form *k* clusters that are “pseudo OTUs” (groupings of related reads). Then, we can assign each query sample’s reads to those “pseudo OTUs” based on distance. A classifier, such as Random Forest, can then be trained to make sample-level predictions on a “pseudo OTU” table made up of the “pseudo OTU” abundance, as defined above, in all samples. Both embedding-based methods learn the sample phenotype by training on each individual read (“read-level”) and on all reads (“sample-level”) rather than read-level-only learning, as for baseline majority vote.

#### Majority vote

The `Read2Pheno` classifier produces a vector of likelihood scores which, given a read, sum to one across all phenotype classes. To get the sample-level prediction, all reads from a sample of interest are classified by `Read2Pheno` model, and the resultant scores are then aggregated by the sample-level predictor. Using body site prediction as an example, there are 5 different body site classes: feces, tongue, skin of hand, skin of head and nostril. We show the diagram of our sample-level predictor in Appendix Sample-level predictor: Majority vote method. Given a sample of interest, the reads associated with this sample are first predicted by `Read2Pheno` classifier. Notice that some species can be found in multiple body sites. Therefore, performing a hard call on which body sites a read originates from can be misleading. To alleviate this problem, if needed, the sample-level predictor contains a read caller function that can assign one read to multiple body sites by applying a threshold to the output of `Read2Pheno` for the read. In our implementation, if the likelihood score of the read from a body site is greater than chance (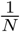, where *N* is the number of body sites in the training data), the vote count of that particular body site will increment by 1 (see the “Read Abundance” block in Appendix Sample-level predictor: Majority vote method). For example, suppose there are three target body sites: skin (i.e., dermal samples), gut (i.e., fecal samples), and tongue (i.e., oral samples). If a read were predicted to be from gut, skin and oral samples with scores of 0.51, 0.43 and 0.06 respectively, both the vote counts of feces class and skin class would increment by 1 (since the likelihood of these two body sites are greater than 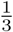). Finally, once all reads have been counted, the `argmax` of vote count vector is taken to predict the sample-level body site.

#### Sample-level embedding

The attention layer of the `Read2Pheno` classifier produces a *N*_*h*_-dimensional embedded vector (see Figure 1) that is a meaningful numerical representation of each 16S rRNA read. For sample-level classification, we first use the trained `Read2Pheno` model to encode all reads per sample into the *N*_*h*_-dimensional vectors. Then, we average the read vectors to form a sample-level embedding. We can then train a classifier (e.g. Random Forest) on the sample-level embeddings to predict phenotype. We show the training and testing process of such method in Appendix Sample-level predictor: Sample-level embedding.

#### “Pseudo OTU” table

While both embedding based sample prediction methods begin with read-level embeddings of the training data, they differ in sample-level training. Instead of taking the average of the trained read embeddings, we use a *k-means* algorithm to cluster the read embeddings of training data into 1000 clusters [38]. Then, all reads in each query sample can be assigned to those clusters. Effectively, the clusters represent related sequences, which are called “Pseudo OTUs”. We compute the number of reads assigned to each pseudo OTU for each sample to create a “Pseudo OTU” table: a matrix of pseudo OTUs vs. samples. Like regular OTU tables, the Pseudo OTU table can train a classifier, such as Random Forest, for sample-level phenotype prediction. A diagram of this process is available in Appendix Sample-level predictor: Pseudo-OTU.

### Data Preparation for Model Evaluation

#### American Gut Project (AGP) dataset

The AGP dataset used for model evaluation in this paper is a subset of data from the American Gut Project [49]. As of May 2017, the AGP included microbial sequence data from in total 15,096 samples from 11,336 human participants and that number continues to grow as the project is ongoing [49]. We focus on samples from five major body sites (*N*_*y*_ = 5): feces, tongue, skin of hand, skin of head and nostril. As mentioned in American Gut Project’s documentation, some bloomed organisms were contained in samples analyzed early in the American Gut Project because of increased shipping time and delay between when samples were collected and when they were put on ice. As a result, bloom sequences should be removed in preprocessing process by American Gut Project. In this paper, we use the latest filtered sequences and OTU table deposited in ftp://ftp.microbio.me/AmericanGut/latest as of 2018/12. All reads have been trimmed to 100 base pairs, so that *T* `=` 100 in Figure 1.

#### Gevers dataset

The Gevers dataset used for model evaluation in this paper is a subset of an inflammatory bowel disease (IBD) dataset [7] (NCBI SRA index: PRJNA237362 in NBCI). Sample metadata label them as being IBD or Non-IBD (*N*_*y*_ = 2). Here, we refer to IBD samples as “CD” (Crohn’s Disease), and the Non-IBD ones as “Not IBD” (disease-negative). We merge paired reads using QIIME [53] and trim them to 160 base pairs (i.e., with the first 10 removed, the following 160 base pairs kept and the rest discarded), so that *T* = 160 in Figure 1.

#### Experimental setup for American Gut Project dataset and Gevers dataset

First, we filter out samples with less than 10,000 reads. Then, we randomly select 161 samples from American Gut Project dataset and 221 samples for Gevers dataset per class as our experimental dataset to balance the class distribution (resulting in total 805 samples in AGP experimental dataset and 442 samples in the Gevers experimental dataset). The number of samples are selected based on the least number of sample per class after filtering for each dataset. Next, we randomly select a certain number of samples per class for training and leave out the rest for testing. For the AGP dataset, 10, 80 and 150 samples per class are randomly selected for training (resulting in 50, 400 and 750 samples total respectively). For the Gevers dataset, 20, 80 and 200 samples per class are randomly selected as training data (resulting in 40, 160 and 400 samples total respectively).

For the AGP dataset-based experiment used for attention interpretation, we randomly select 10 samples per class for training, resulting, in total, 50 samples and 1,503,639 reads for training. The rest of the samples form the testing dataset. We randomly select 10 samples per class as the candidate visualization set. For the Gevers dataset-based experiment used for attention interpretation, we select 40 samples (20 from the IBD class and 20 from non-IBD) by random and collect 1,678,464 reads for training (around 42,000 reads per sample). The remaining samples (442 - #_of_training) are used for testing. We again randomly select 10 samples per class from the testing dataset as the candidate visualization set. After we select the candidate visualization set for both attention interpretation experiments, we use the QIIME [53] implementation of the Ribosomal Database Project (RDP) [54] taxonomic classification to assign the genus-level labels to reads in the candidate visualization set. Then, reads with less than a 80% RDP confidence score on genus level are removed from the visualization set. Finally, in order to efficiently extract intermediate layer outputs and generate visualizations, an arbitrary subset, 100,000 reads from the qualified visualization set, are randomly sampled for the final visualization and interpretation. All reads in the final visualization set have a genus-level label and phenotype (i.e., body site or disease diagnosis) label. For the AGP visualization set, we further merge the skin-associated label, namely, skin of head, skin of hand and nostril into one single skin class to simplify the visualization. As a result, the visualization set reflects 3 body site classifications instead of 5. We use the experimental setup for American Gut dataset as an example to show the overall training and testing experiment in Appendix Overall training and testing experiment.

#### SILVA dataset and experimental setup

The SILVA 16S taxonomic QIIME-compatible dataset is used to construct our experimental dataset [50, 51]. There are 369,953 sequences total in the original dataset. Among those sequences, there are 268,225 which have a genus-level label, and those sequences come from 6,618 genera. We select the genera that have over 100 representative sequences and collected all sequences from these genera to form our experimental dataset. We thereby include in total 204,789 sequences from *N*_*y*_ = 495 genera in our experimental dataset. Our dataset covers around 76.35% sequences and 7.48% of the genera in original dataset. Sequences are first one-hot coded according to the map shown in Appendix One-hot coding map. Then, we right pad the sequences with zero vectors to the nearest hundred and grouped sequences based on the padded length (resulting 11 groups that have 100bp increment size in the range of *T* = 900 → 1900). For example, a sequence of length 1001 will be padded with 99 zero vectors to a total length of 1100bp. Then, those sequences are stored in the same matrix per length group. In this way, model can be trained by sequences with similar length at a time to improve the training efficiency. We then randomly split the dataset into 80% sequences as training and 20% as testing.

### The Read2Pheno Training Process

We train the `Read2Pheno` model with reads from training dataset, preprocessed and selected as described above, labeling reads by sample phenotype. Since the `Read2Pheno` model should be trained and optimized for read level prediction, sample-level predictors are trained separately after the `Read2Pheno` model training is completed. For each testing sample, all reads are classified by the `Read2Pheno` model. We then aggregate the read level information encoded by the `Read2Pheno` model using methods described in Section Sample-level predictor to make sample-level predictions. We show a schematic of the training process in Appendix `Read2Pheno` training process.

We randomly sample an equal number of samples from each class to form the training set. Then, we label all reads associated with those samples by their sample-level label and shuffled. All reads are one-hot coded according to the coding map in Appendix One-hot coding map. Then, the data is fed to the `Read2Pheno` model for training. The reason we train our neural network in read level instead of sample level is two-fold: 1) our read level model can highlight informative regions in each input sequence; 2) there are relative less number of examples to train a complex neural network model in sample level than in read level. As discussed in Section Model selection on American Gut dataset, we further show that the read level model trained with a dozen of samples performs comparably to read level model trained with hundreds of samples.

Our deep learning model is implemented in Keras (version 2.2.2) with Tensorflow (version 1.9.0) backend. If the number of classes is greater than 2, then categorical cross-entropy can be used as the loss function. Otherwise, binary cross-entropy is the recommended loss function. Adam optimization with default setting and a learning rate of 0.001 is used to train the model. The model was trained and evaluated on the Extreme Science and Engineering Discovery Environment (XSEDE) [55] for 10 epochs. We also made a python module of the `Read2Pheno` model in Github https://github.com/EESI/sequence_attention.

### Model Interpretation and Read Visualization

The `Read2Pheno` model has an LSTM layer; consequently, sequential information are encoded and circulated in hidden states. The intermediate output, labeled as “Read embedding” in Figure 1 is a *N*_*h*_-dimensional vector. This read embedding vector is an average of hidden states across all bases weighted by the attention weights, labeled as “Attention Weights” in Figure 1. The *N*_*h*_-dimensional embedding vector can be considered as a numerical representation of the input DNA/RNA read. Therefore, similar reads should be embedded to vectors that are close to each other in *N*_*h*_-dimensional space, whereas differing reads should be embedded far away from each other. This type of relationship may be shown by plotting the *N*_*h*_-dimensional vector representations of the reads in a 2-dimensional space. Accordingly, we use Principle Component Analysis (PCA) to reduce the dimensionality of all reads in visualization set to 2-dimension by projecting them onto the top 2 principle components that explain the most variation.

Inspired by “WebLogo” [56, 57], we also use a “sequence logo” to visualize significant features contained by the sequence. The reads from same genus are similar to each other, with mutations at certain positions. We thus group the visualization reads from the same genus together for further exploratory visualization. In this study, we use QIIME [53] implementation of RDP [54] taxonomic classification method to predict the genus level label for our visualization reads.

We calculate the overall Shannon Entropy of a group of reads (reads from a genus) by:

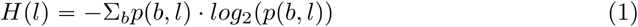

where *b* is the nucleotide base, *b* ∈ {*A, T, G, C*}, *l* is the position of the sequence, *l* ∈ (0, length(seq)]. *p* (*b, l*) can be estimated by *f* (*b, l*), the normalized nucleotide frequency of base *b* at position *l*. The sequence logo for a given phenotype class can be calculated by Equation 2 where *c* is the phenotype label and *f*_*c*_ (*b, l*) is the normalized nucleotide frequency of base *b* at position *l* among reads from phenotype *c*.

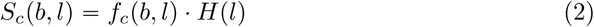

Positions with high nucleotide variance will have high entropy and therefore the sequence logo is a good measure of “importance” (variation) of nucleotides, which we use as our baseline importance measure.

As presented in Figure 1, a dense attention layer learns to predict the importance of bases by hidden states output from Bi-LSTM layer. The output of the attention layer, “Attention Weights”, is a vector of input read length, *T*, wherein each value represents the importance of the hidden state corresponding to said position. This vector will indicate what region of the input sequence the model has been found to be most informative. Therefore, we use the attention weights for the input reads as the model’s predicted importance measure. Among reads from the same genus, the attention weights for reads from the same phenotype are averaged. The mean attention weight vector highlights the informative sequence regions for a phenotype for this genus. The attention measure is thus defined by Equation 3.

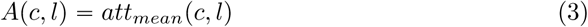

where *att*_*mean*_ (*c, l*) is the mean attention weight of reads from a phenotype *c* at position *l*.

## Results

As described in detail below, we analyzed three distinct 16s rRNA amplicon sequence data sets: 1) data provided by the American Gut Project (AGP), in which samples are labeled by body site origin and thereby reflect microbiome phenotype (i.e., properties of a microbial community); 2) data published by Gevers *et al*. (Gevers), which is labeled by disease diagnosis, i.e., host phenotype; and 3) the SILVA rRNA database, large corpus of comprehensive and quality checked 16s rRNA sequence dataset with taxonomic labels. Our goals for each type of data set were to evaluate 1) the performance of attention-based deep learning models at predicting phenotype and taxonomy as compared to existing baseline methods, and 2) interpretability gains afforded by intermediate layer outputs of attention-based deep neural networks through visualizing the ordination of sequence embedding vectors and informative regions of sequences highlighted by attention weights.

### Microbiome Phenotype (Body Site) Prediction based on American Gut Project (AGP) Data

We evaluated our proposed `Read2Pheno` attention model on a subset of the American Gut Project (AGP) dataset. The AGP dataset contains sequencing data from the largest crowd-sourced citizen science project to date [49]. Our experimental dataset contains 805 samples obtained from five body sites: feces, tongue, skin of hand, skin of head, and nostril.

#### Model selection on American Gut dataset

We use the training data of 50 samples to perform a 5-fold cross validation to fine tune the hyperparameters of our model. The hyperparameter search space can be found in Appendix Hyperparameter search space table. The 5-fold cross validation yielded *N*_*c*_ = 256 filters in CNN layers, *N*_*h*_ = 64 units in LSTM layer, dropout rate of 0, and learning rate of 0.001 as the hyperparameters as producing the best read level classification accuracy on the training dataset. Accordingly, we incorporated these hyperparameters in our model. We also performed the same hyperparameter sweep process on other models with related architectures: the Bi-LSTM model,

Attention-based Bi-LSTM model, CNN model, Attention-based CNN model, CNN-Bi-LSTM model and Attention-based CNN-Bi-LSTM model. We use the same architecture for CNN and Bi-LSTM layers in the models described in Figure 1. Table 1 shows the best set of hyperparameters for all 6 models. In Table 1, the CNN column shows the optimal number of convolutional filters, *N*_*c*_. The RNN column shows the optimal number of hidden nodes in Bi-LSTM, *N*_*h*_. DP refers to the dropout rate (probability of training to a particular hidden node in the layer) and LR is the learning rate (amount weights are updated in each step) used in Adam optimizer. From the table, classifiers constructed with only a Bi-LSTM layer or a CNN layer have suboptimal accuracy compared to more complex models. With the help of an attention mechanism, the CNN model achieves better accuracy, but the Bi-LSTM model doesn’t benefit from the attention layer. The classifier which combines CNN layers, a Bi-LSTM layer and an attention layer results in the best accuracy classification following 5-fold cross-validation. Although the model without an attention layer achieves a similar accuracy, the interpretability of the attention-based model is superior, as shown in the following section discussing sample-level prediction. To evaluate the effect of using small number of training samples for `Read2Pheno` classifier training. We design an independent experiment: we first hold out 55 samples as testing, then we train the `Read2Pheno` model with reads from 5, 25, 50, 100, 500 and 750 samples from the rest of samples and evaluate the sample level performance of those models by the 55 held-out testing set (here we use the sample-level embedding method for sample prediction). For sample-level phenotype prediction, there are two types of ways to train the Random Forest (RF) model: 1) we use the exactly same training set to train RF as used to train `Read2Pheno` classifier and then test on the test set; 2) despite of the number of samples used to train `Read2Pheno` classifier, we use all 750 samples in training set train the RF and test on the test set. We show the performance (The blue line shows the training type 1 and the orange line shows the training type 2) in Appendix Training data size effect of `Read2Pheno` classifier. The blue curve shows that as more samples used for training, the sample-level accuracy increases. The orange curve shows that although `Read2Pheno` classifiers are trained with different samples, as long as the sample-level prediction model is trained with more samples, the performance is pretty stable. This indicates that the `Read2Pheno` classifier can learn a meaningful embedding with only a small number of samples. In fact, there are usually a great number of reads in a few samples. For example, in AGP dataset, there are over 1 million reads in 50 samples). Therefore, for further downstreaming analysis including embedding visualization and attention weights visualization, we use the model train by 50 samples.

**Table 1.** Training Accuracy Comparison. Results of 5-fold cross validation for model/hyperparameter selection

| Model | CNN ($N_c$) | RNN ($N_h$) | DP | LR | Acc (± Std) |
| --- | --- | --- | --- | --- | --- |
| Bi-LSTM | - | 128 | 0.25 | 0.005 | 0.734 ($\pm 0.002$) |
| Bi-LSTM+ATT | - | 128 | 0.25 | 0.005 | 0.732 ($\pm 0.003$) |
| CNN | 128 | - | 0.25 | 0.001 | 0.738 ($\pm 0.001$) |
| CNN+ATT | 256 | - | 0.25 | 0.001 | 0.740 ($\pm 0.001$) |
| CNN+Bi-LSTM | 256 | 128 | 0.25 | 0.001 | 0.742 ($\pm 0.001$) |
| CNN+Bi-LSTM+ATT | 256 | 64 | 0 | 0.001 | 0.742 ($\pm 0.001$) |

#### Sample-level phenotype prediction

Sample-level phenotypes are predicted by sample-level predictors as described in the Methods section (in Section Sample-level predictor). Table 2 compares the accuracy of our deep learning approaches (a `Read2Pheno` model trained for 10 epochs with various sample-level classification strategies) against Random Forest (RF) baseline approaches, which are trained on three different types of features: *k*-mer counts, OTU tables and ASV tables generated by Dada2 [58]. In Table 2, we compare the training dataset size’s effect on the models’ accuracies (all `Read2Pheno` models are trained for 10 epochs and followed by various sample-level classification strategies for sample level predictions). For example, for the training set of size of 50 samples, our method was trained for 10 epochs with 50 samples and tested by 755 samples, whereas a RF classifier with 100 estimators was trained by the same 50 training samples and tested by 755 samples using the 9-mer frequency feature table, OTU table and ASV table respectively. We use a 9-mer frequency feature table because the filter window size of our convolutional block is 9. As expected, adding more training data increases performance. While training an RF model on raw 9-mers performs very well for all training sizes, our sample embedding and pseudo-OTU methods outperform the 97% identity OTU tables. Moreover, prediction accuracies using the pseudo-OTU approach are competitive with using 9-mer raw features. But accuracy is not the ultimate objective of these phenotype prediction experiments. Our read-level and sample-level embeddings can be interpreted to visualize read-phenotype and read-taxa relationships, a task that 9-mer features cannot accomplish alone. We hypothesize that our embedding approach is able to perform well at clustering sequences from similar taxa together. The target classes in this comparison are 5 body sites: feces, tongue, skin of hand, skin of head and nostril. Unlike the pseudo-OTU method, an OTU table-based method can only identify informative OTUs, rather than informative sequence context (like Figure 3). Moreover, *k*-mer based methods can highlight useful local *k*-mer information, but cannot easily interpret informative taxa (like Figure 2) and sequential order information. By contrast, our attention-based model can provide deeper feature visualizations (Figure 2, 3 and Appendix 2D Visualization of *Prevotella* reads) and interpretation.

**Table 2.** American Gut Data Testing Accuracy Comparison. Unlike sample-level classification methods that use OTU/ASV tables and /*k*-mers (e.g. 9-mers) as features, our proposed model is trained on reads. Then, the read-level results are fused by the sample-level predictor using three methods as described in this paper. By increasing the number of samples in the training data, we compare the read-level classifier’s ability to learn sample-level predictive taxa/information from limited data sizes. Accuracies are averaged and standard deviation is measured over 5 randomly selected data with replacement experiments. We show sample-level prediction for the proposed methods are competitive with prediction from OTU tables and will allow interpretable representations shown in subsequent sections.

|  |  | Training Set Size |  |  |
| --- | --- | --- | --- | --- |
| Category | Method | 50 | 400 | 750 |
| Traditional | 9-mer table | 0.808<br>(± 0.039) | 0.887<br>(± 0.009) | 0.903<br>(± 0.024) |
| | OTU table | 0.731<br>(± 0.021) | 0.816<br>(± 0.010) | 0.831<br>(± 0.030) |
| | ASV table | 0.769<br>(± 0.034) | 0.840<br>(± 0.014) | 0.866<br>(± 0.024) |
| Proposed | Majority vote | 0.730<br>(± 0.040) | 0.794<br>(± 0.012) | 0.795<br>(± 0.030) |
| | Sample embedding | 0.751<br>(± 0.012) | 0.816<br>(± 0.010) | 0.821<br>(± 0.025) |
| | Pseudo OTU | 0.784<br>(± 0.039) | 0.858<br>(± 0.013) | 0.881<br>(± 0.026) |

**Figure 2.**
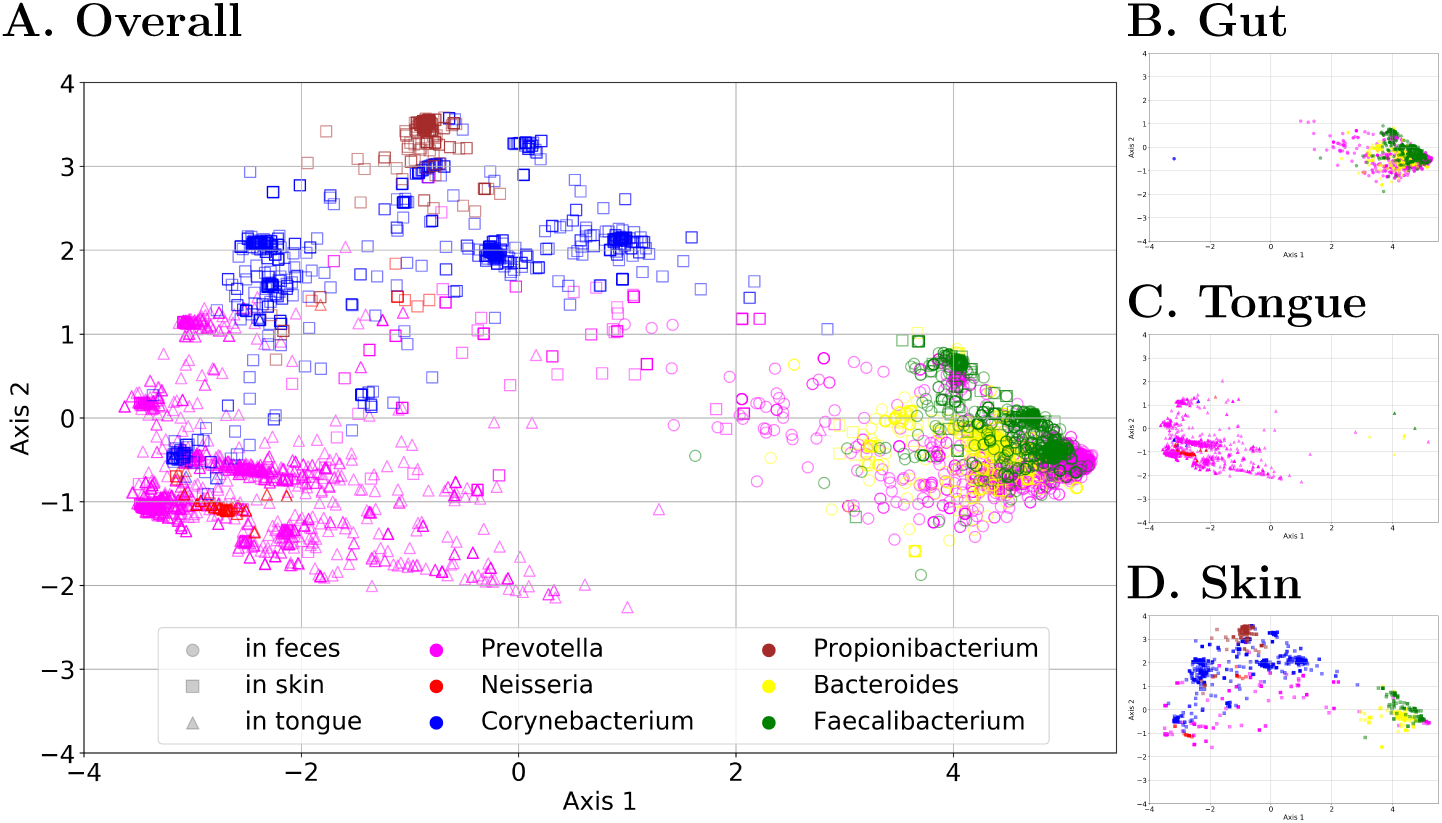
2-D projection of embedded read vectors from all body sites (A), gut (B), tongue (C) and skin (D). ‘□’s are reads from the skin, ‘○’ s are reads from the tongue and ‘Δ’s are reads from the gut/stool. The neural network is learning the 16S rRNA gene association to taxonomy and body site without the access to taxonomy label of the reads.

**Figure 3.**
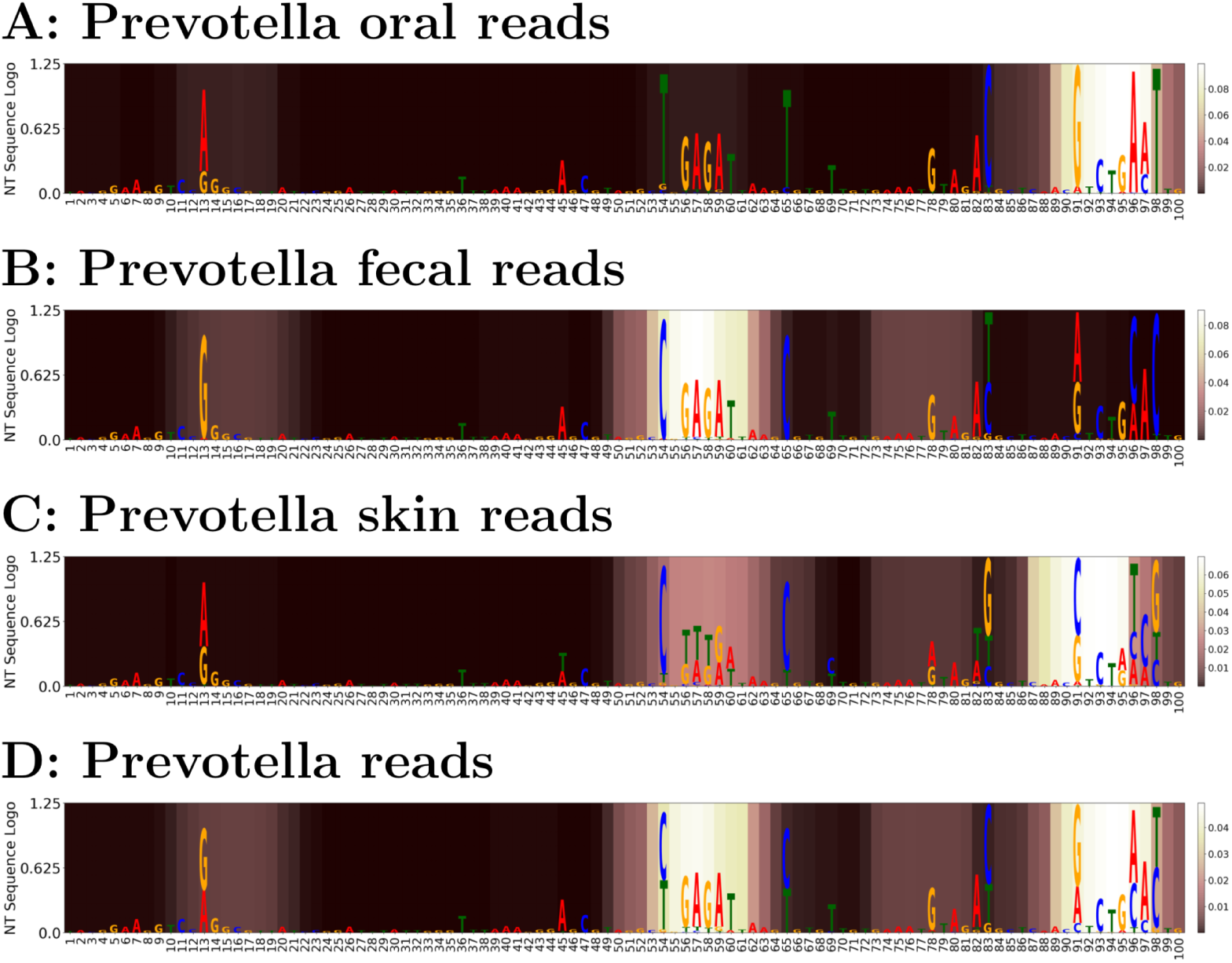
Comparison between average *Prevotella* reads attention and nucleotide frequency entropy in form of nucleotide sequence logo. A: oral reads; B: fecal reads; C: skin reads; D: overall attention. In each body site, nucleotide frequencies are scaled by the overall entropy for all *Prevotella* testing reads and plotted as a sequence logo, with average attention weights represented by a color map where lighter background shading represents larger values for attention weights, in contrast with darker background shading for smaller attention weights.

#### Read embedding visualization

To illustrate how the model can learn taxonomic classes despite only having phenotype labels, we visualize the embedded vectors for reads from 6 selected genera.

Figure 2 shows the 2-D principal component analysis (PCA) projection of embedded vectors of 16S rRNA reads (the intermediate output vectors of the Multiplication Layer in Figure 1) of 6 genera from the American Gut data. Each point represents a read, in which the color represents the genus label (determined by RDP [54]) and marker shape represents the original body site (as determined from the body site label). As mentioned in Section Experimental setup for American Gut Project dataset and Gevers dataset, to produce a clear visualization, we merge skin of hand, skin of head and nostril to one single skin class. In Figure 2, reads from one body site are clustered closer together than to other body sites in the embedding space. For example, as shown in Figure 2B, reads from gut are embedded together. In addition,, most of the reads from particular genera are closely embedded together. This illustrates that even though the model is not optimized for taxonomic classification, the neural network is still learning the 16s rRNA variable V4 region—which contains mutations that indicate different taxa—of the input reads in the embedding space. Notably, for *Prevotella*, most fecal-associated reads separate from the oral-associated ones, demonstrating that the model can discern sub-genera. It is most likely that within these sub-genera, different species have preference for different body sites. This kind of intra-genus separation does not appear for all genera, however. This is to be expected, since the same 16S rRNA read may exist within multiple body sites, which would make it hard for the model to predict such a read correctly. Nevertheless, some skin-associated *Corynebacterium* strains separate out, revealing which intra-genus 16s rRNA variants can and cannot be learned.

#### *Prevotella* Case Study

To understand which features facilitate class separation, we again inspected the read embedded vectors for genera which separated well for the body site isolation source. Notice in Figure 2, reads from *Prevotella* formed two major clusters corresponding to two body sites, namely, tongue and feces. Therefore, we analyze *Prevotella* as an exemplary demonstration of the interpretability of the attention learning mechanism.

As shown in Appendix 2D Visualization of *Prevotella* reads, a 2-D PCA projection of embedded vectors from *Prevotella* test reads forms two well-separated clusters, which correspond to tongue and feces. The *Prevotella* test reads were classified to the correct body site source with 91.31% accuracy. To visualize the regions that are most informative to this classification, Figure 3 shows which high entropy positions also have high attention using the method described in sample-level predictors section. Panel D of Fig 3 shows that the middle and end portions of the 100 bp trimmed reads are most important for phenotype classification, with the former playing a more important role in distinguishing fecal reads (panel B) while the latter is more important for oral and skin reads (panels A and C). For visualization, the attention weights are smoothed by a moving average of window size of 9 (i.e., the size used in the convolutional filter of the model).

Inspecting the output of the attention layer for *Prevotella*, we can see which areas of the 16s rRNA V4 region that the network is paying attention to (by the brightness of the highlight) to make this classification, shown in Figure 3. We can see that the end of ths 16S V4 region is the most important for identifying oral and skin reads, while the middle region is the most important for gut/fecal reads. However, there are slight differences – for example, an area at the beginning of the V4 region has some importance to also help identify the gut, and to some extent oral, reads—as opposed to getting no attention weighting for skin reads. When comparing oral and skin reads, the middle region is the second most important to identify skin reads. This region may help resolve oral/skin body sites that have similar nucleotides at the end of the reads.

For comparison, the entropy of the sequences within the Prevotella body site combinations are shown (with the whole general attention shown in Figure 3D). We can see that the attention model is generally learning areas of the variable region that have high entropy. However, it is also learning slight differences between the signatures of these regions. For example, both gut/skin reads tend to have C’s located at positions 54 and 65 while gut/oral reads tend to have `GAGA` at 56 → 59. Thus, the particular combination of `C-GAGA-----C` is unique to gut reads and therefore, a high attention weight is placed on this region to distinguish gut reads from other body sites. In sum, the attention weights shown in Figure 3 will reflect nucleotide variation found in training sequence data, which, in turn, helps the model predict body site labels from the input reads.

### Host Phenotype (Clinical Diagnosis) Prediction based on Gevers Inflammatory Bowel Disease Data

We further evaluate `Read2Pheno` performance on a distinct set of sequence data, the Gevers dataset, which as described in Section Data Preparation for Model Evaluation, is a subset of data from an inflammatory bowel disease (IBD) study in which samples were identified as being from patients who were diagnosed with inflammatory bowel disease (IBD) and not [7].

#### Sample-level phenotype prediction

Sample-level phenotype prediction is accomplished by 1) the sample-level predictors discussed in Appendix Sample-level predictor: Majority vote method with Read Caller threshold of 0.5 (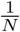, where *N* = 2 for two classes); 2) sample-level embedding based Random Forest; 3) “Pseudo OTU” table based on a Random Forest trained on 1000 Pseudo OTUs. Table 3 compares the accuracy of our model against three baseline methods: Random Forests trained on (1) *k*-mers, (2) OTU tables, and (3) amplicon sequence variants (via Dada2). We show the testing accuracy for different training data sizes in Table 3. For example, for the training set size of 40 samples, we trained our method for 10 epochs, and compared that to training a Random Forest classifier with 100 estimators using the various baseline methods. We compare against a Random Forest trained on the 9-mer frequency feature table. *K* 9 seems like a reasonable choice because the filter window size of our convolutional block is 9. The target classes in this comparison are 2 states: IBD and Non-IBD. From the table, we can see the performance of our “Pseudo OTU” based method is the comparable to competing methods. When trained with 40 samples and tested on the rest, 9-mer table based model works the best and our Pseudo OTU model is comparable to the OTU table based method. As more samples are used for training, the performances of all models are improved. In general, among our proposed models, the Pseudo OTU model consistently works the best. The Pseudo OTU model is often comparable to OTU and ASV based model but slightly underperforms 9-mer table based model.

**Table 3.** Gevers Dataset Testing Accuracy Comparison. Unlike OTU/*k*-mer based classifiers, which are trained at sample-level, our proposed model is trained at the read level before read level results are then fused by sample-level predictor. This comparison, over 40, 160, and 400 samples in the training data shows that the read level classifier learns predictive taxa/information and the sample-level prediction for the proposed methods are competitive with prediction from OTU tables and will allow interpretable representations shown in the subsequent sections. The obtained accuracy values are averaged, and the standard deviation is computed, over 5 experiments in which we randomly selected training-testing data splits with replacement.

|  |  | Training Set Size |  |  |
| --- | --- | --- | --- | --- |
| Category | Method | 40 | 160 | 400 |
| Traditional | 9-mer table | 0.715<br>($\pm 0.029$) | 0.787<br>($\pm 0.030$) | 0.848<br>($\pm 0.055$) |
| | OTU table | 0.684<br>($\pm 0.016$) | 0.768<br>($\pm 0.018$) | 0.843<br>($\pm 0.0548$) |
| | ASV table | 0.669<br>($\pm 0.019$) | 0.765<br>($\pm 0.022$) | 0.819<br>($\pm 0.078$) |
| Proposed | Majority vote | 0.653<br>($\pm 0.043$) | 0.690<br>($\pm 0.023$) | 0.729<br>($\pm 0.062$) |
| | Sample embedding | 0.650<br>($\pm 0.016$) | 0.726<br>($\pm 0.029$) | 0.762<br>($\pm 0.069$) |
| | Pseudo OTU | 0.689<br>($\pm 0.031$) | 0.779<br>($\pm 0.014$) | 0.833<br>($\pm 0.058$) |

#### Read embedding visualization

To inspect how the read embeddings identified by `Read2Pheno` perform, here we visualize the embedded vectors for reads from 4 selected genera (*Blautia, Roseburia, Ruminococcus*, and *Pseudomonas*). Appendix 2-D projection of embedded read vectors for Gevers dataset shows the 2-D PCA projection of embedded vectors (the intermediate output vectors of the multiplication layer in Figure 1) of a selection of 4 genera from 16S rRNA reads from the Gevers Crohn’s disease dataset [7]. Each point represents a read, in which the color represents the genus label (determined by RDP [54]) and marker shape represents the disease state (CD: Crohn’s disease; Not IBD: No inflammatory bowel disease diagnosis). In Appendix 2-D projection of embedded read vectors for Gevers dataset, most of the time, reads from one genus are closely embedded together. However, the Not IBD samples for the *Roseburia* and *Ruminococcus* genera have the widest spread in PCA. In fact, we can see multiple clusters in most of the genera, suggesting that different sub-genera cluster together and can be associated with different phenotypes. In addition, reads identified as disease-positive (“CD”) are generally clustered in lower right of the figure, while reads labeled as disease-negative (“Not IBD”) are clustered in upper left of the figure. Even though the model has not been trained to do taxonomic classification—only disease phenotype—it can still reflect what it has learned from the sequence structure of the input reads in the embedding space to reveal taxonomic structure.

Indeed, within *Blautia* and *Ruminococcus* genera, there are at least one cluster corresponding to disease (“CD”) and another cluster corresponding to non-disease (“Not IBD”). This further demonstrates that the `Read2Pheno` model can discern sub-genera that have associations with different phenotypes. Finally, most *Pseudomonas* reads are disease-positive (labeled as “CD”), and they are also embedded in lower right corner in the figure, which shows that the `Read2Pheno` model has predicted that those reads are disease-positive. This is consistent with the association between *Pseudomonas* species and IBD, which has been described extensively in the literature [59–61]. We further show the interpretability of our model in Section Case studies of Gevers dataset.

### Taxonomy Prediction based on SILVA Full Length 16S rRNA Sequence Data

We further evaluate the capability of our proposed `Read2Pheno` model to analyze and learn which regions of the full-length 16S ribosomal RNA sequence are useful for predicting the genus level taxonomic label. The experimental dataset in this section is constructed from the SILVA 16S ribosomal RNA gene database [50] and their manually curated taxonomy [51] (i.e., Release 132 16S sequences with 99% identity criterion to remove highly identical sequences).

#### Full length 16S rRNA taxonomic classification

Unlike the other results presented above, here, we train our proposed `Read2Pheno` model on *taxonomic* classification specifically. In particular, we train the model on the SILVA training dataset for 40 epochs. We adopt the same set of parameters and NN model architecture used in the previous experiments (256 filters in CNN layers, 64 units in LSTM layer, 0 dropout rate and 0.001 learning rate), except for the number of neurons in output node, *N*_*y*_, which must be set to 495 to accommodate all the genera classes. The same training data is used to train the QIIME [53] implementation of RDP [54] taxonomic classifier. Then, both the RDP classifier and our `Read2Pheno` model are tested by the testing dataset. Table 4 shows the results of both models.

**Table 4.** Accuracy Comparison on the SILVA dataset over 5-fold cross-validation. The proposed model’s performance is slightly below but still competitive to RDP’s accuracy.

| Method | Avg. Accuracy (std) |
| --- | --- |
| RDP implemented in QIIME | 0.976 (± 0.001) |
| Proposed model | 0.959 (± 0.006) |

#### Full length 16S rRNA sequence visualization

We visualize the embedded vectors for full-length 16S rRNA sequences from 7 selected genera in the *Bacillaceae* family in Figure 4. A 2-D principal component analysis (PCA) projects the embedded vectors (the intermediate output vectors of the Multiplication Layer in Figure 1) of the sequences from the selected 7 genera. Each point represents a sequence, in which the color represents the genus label. In Figure 4, most of the sequences from one genus are closely embedded together and sequences from different genera are embedded apart from each other. This illustrates that the model is learning the taxonomic information from the labels, whereas in the American Gut section, taxonomic information was being learned indirectly from phenotype labels. We can also see (although distorted from the 2-D projection) that some *Bacilli* have 16S rRNAs that may be similar to other types of *Bacilli* genera like *Virgi-* and *Oceano-bacillus* but be more distinct from *Geobacillus*. This could indicate misclassifications of these sequences in the standard taxonomy (e.g. Bergey’s Manual of Systematic Bacteriology) or simply evolutionary relatedness between taxa.

**Figure 4.**
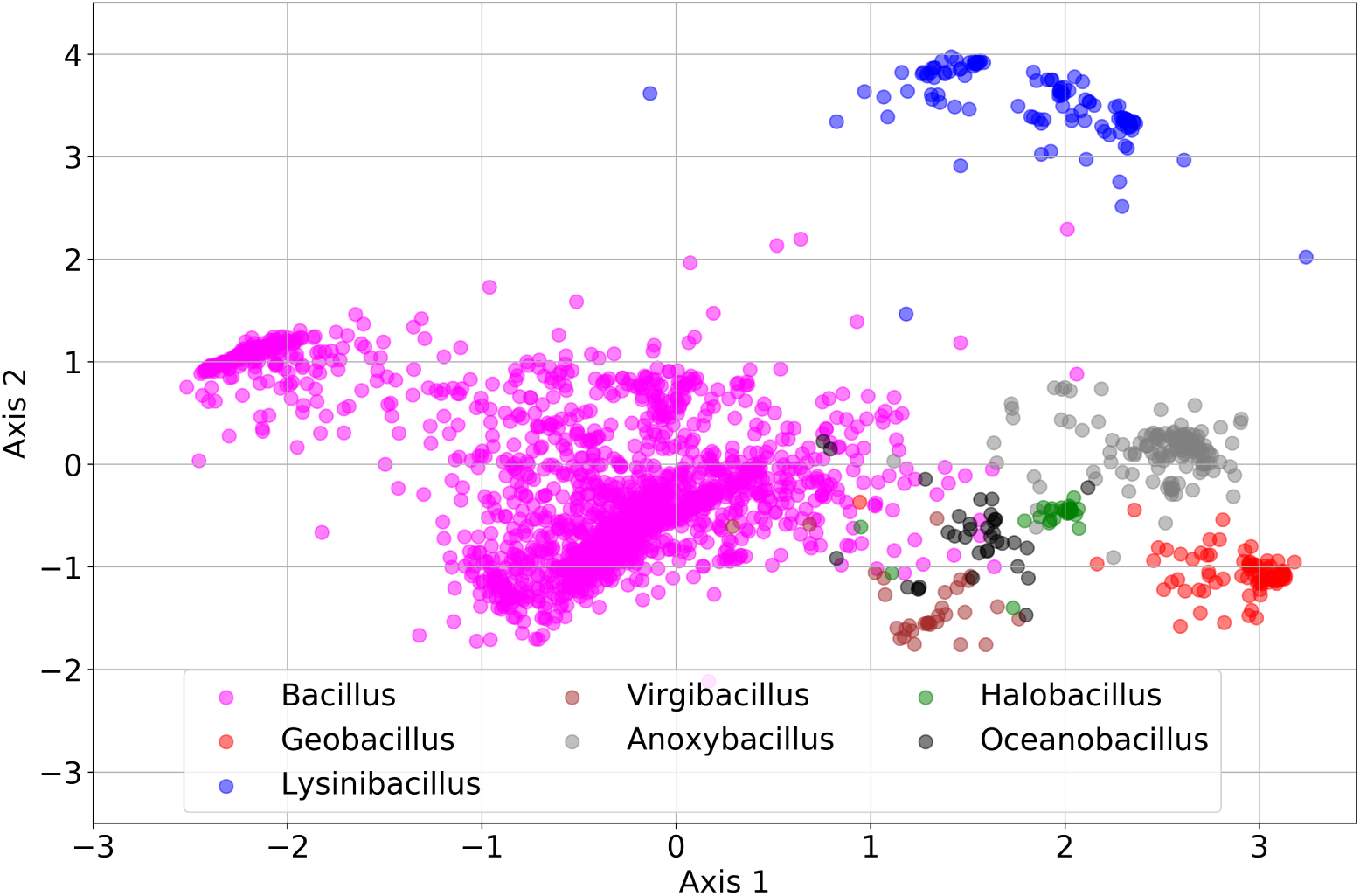
2-D projection of embedded sequence vectors. The model can separate sequences with respect to their genus level label based on their genomic content.

#### Variable Regions discovered by Attention Weights: *Pseudomonas* and *Enterobacter* examples

In Figure 5 and Appendix Average attention weights of *Enterobacter* testing sequences, we visualize the average attention weights of testing sequences from two select genera. For *Pseudomonas* in Figure 5, the top figure shows the averaged attention weights per all testing reads without alignment. The variable regions are labeled according to [62]. Here, different colors correspond to different variable region (from V1 to V9 as shown in the colorbar). As we can see, the attention weights pay attention to nucleotides concentrated on V2 and V3 regions. There are insertion and deletions in different *Pseudomonas* sequences. As a result, the location of a certain context that gain high attention weights can be shifted in different sequences. We applied multiple sequence alignment to align the testing sequences for *Pseudomonas* using the MAFFT on XSEDE [63]. The attention weights are then aligned by the sequence alignment results. Then, the average attention are computed based on these aligned attention vectors. From the bottom figure in Figure 5, we observe that the attention sites narrow down to a few select nucleotide positions despite insertions and deletions in the 16S rRNA evolution. This is evidence that our model is learning specific 16S rRNA nucleotide contexts that are important to the distinction of taxa and decides where to pay the attention based on the context. We further visualize the attention weights of a real *Pseudomonas aeruginosa* sequence provided by [64] in Appendix Secondary structure and attention weights of a *Pseudomonas aeruginosa* sequence. The positions that have an attention weight greater than the mean attention weight cross the whole sequence is highlighted on the secondary structure figure. The attention-highlighted regions coincide with sub-regions of V2, V3, and V4 variable regions determined by [64]. Moreover, we notice that attentions are paid in the junctions in the secondary structure by the model. We conjecture that those junctions are related to molecular interaction and the nucleotides at those positions can contribute to the angle/structure of the arms associated with the junction.

**Figure 5.**
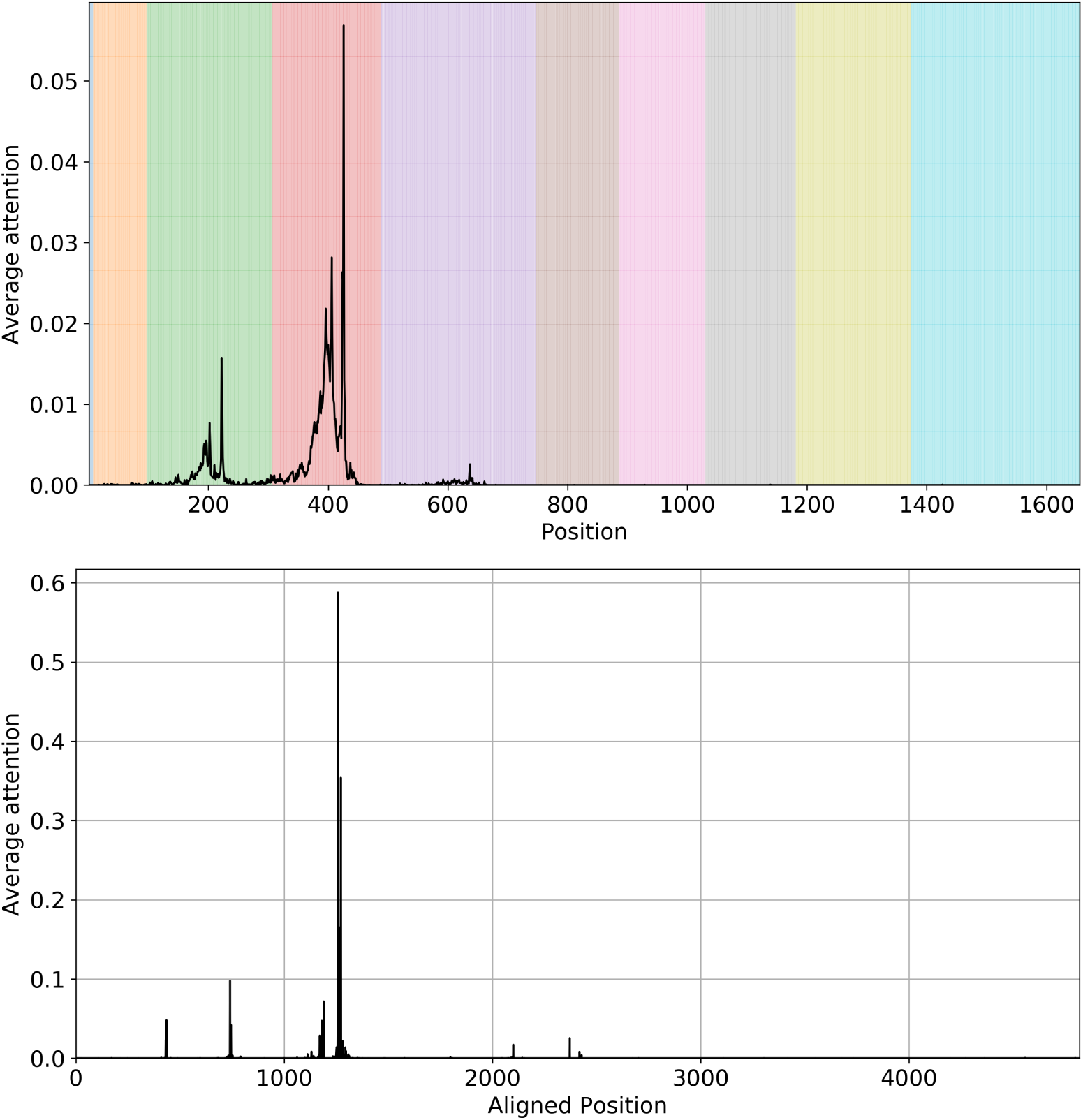
Top: Average attention weights for all strains of *Pseudomonas* without alignment; Bottom: Average of attention weights for all strains with alignment. This figure shows that our model can implicitly learn the multiple sequence alignment since there are very few sites with attention – meaning that despite insertions/deletions, the attention is consistently paid to the most informative bases.

We also applied the same analysis on Enterobacter as a comparison (result shown in Appendix Average attention weights of *Enterobacter* testing sequences). For *Enterobacter*, the attention weights are concentrated on V2 and V4. After multiple sequence alignment, we observe concentration of attention weights into mainly two sites (similar narrowing trend as in *Pseudomonas*), which implies that the model can learn contextual information and is robust to insertions/deletions/mutations. We offer the readers our multiple sequence alignment results in FASTA format and the mean attention weights per position after alignment in Appendix Mean attention weights and multiple sequence alignment results aligned for *Pseudomonas* and *Enterobacter*. To offer insight into what the neural network is learning as the most important variable regions for each genus, we calculated the sum of attention weights of testing sequences per variable region (with variable region locations defined by [62]) for each genus and can be found in Appendix Attention weights on variable regions per genus. There are previous works that aimed to find important/predictive variable regions that can inform taxonomic classification and V2, V3, V4 are considered as “informative” [65–68]. Our model, in most cases, pays most attention to the V3 and V4 regions, which many studies now use as common targets. Genera such as *Buchnera, Erwinia*, and *Gemmata* have higher attention weights on the V2 region.

## Discussion

In this paper, we propose an attention-based deep neural network for read level classification that can reveal informative regions that are relevant to the phenotypic classification (classification of 16S rRNA reads from microbiome to phenotype). We have shown that attention-based deep learning, and specifically our proposed `Read2Pheno` models are capable of comparable accuracy prediction performance while offering automated model interpretation on three distinct kinds of tasks: (1) prediction of microbiome phenotype (i.e., the emergent property of a microbial community), (2) prediction of host phenotype (i.e., clinical disease diagnosis), and (3) taxonomic classification of full length 16S rRNA sequences. The implications of our attention-based deep learning methodology, as implemented and evaluated on these tasks as proof-of-concept, are discussed in further detail below.

As proof-of-concept for microbiome and phenotype prediction, we have focused on two large-scale microbiome datasets. *First*, we have analyzed data from the American Gut Project, which provides a comprehensive open-source and open-access set of human microbiome 16S rRNA samples for scientific use [49, 69]. The recent studies of microbiomes inhabiting sites on the human body (particularly the large intestine) have revealed the complex nature of microbial community interactions [69]. 16S ribosomal RNA is not only useful for identifying organisms using the phylogenetic tree of life, but the phylogenetic branch distance shared between samples serves as a comparative distance metric [70]. Utilizing the AGP’s large collection of samples, where we can identify organisms via 16S rRNA, allows us to begin to understand microbial community dynamics in hosts and the environment [71]. The AGP dataset thus provides real-world data to develop and validate phenotype prediction algorithms. *Second*, to analyze host phenotype, we have further looked to gut data with clinical significance as well. Crohn’s disease (CD), a chronic relapsing inflammatory bowel disease (IBD), is increasing in prevalence worldwide [72]. Researchers have been exploring different methods to predict Crohn’s disease based using microbiome data, for example to identify the microbial taxa that associated with the disease using 16S rRNA survey data [7, 9, 73]. We have further evaluated `Read2Pheno` and sample-level classification using a dataset based on clinical evaluation provided by Gevers *et al*. Developing a better understanding of *the* IBD microbial signature will present a critical step towards improved clinical diagnosis and discovery of a cure.

Our results with these data sets show that we can keep both local information (*k*-mers) and contextual information (sequential order of *k*-mers) of 16S rRNA regions without the need of abundance table such as OTU or ASV for phenotype prediction and achieve comparable performance to phenotype classification based on OTUs/ASVs. The number of samples is relative small to train a complex deep neural network for sample level prediction and can lead to overfitting (especially when the variation in the dataset is low) [74, 75]. For example, there are over 1 million parameters in the proposed deep neural network model for AGP experimental data, we have only 805 samples (perfectly balanced), however, for training. And the number of data points, *n*, should be no less than some multiple (say 5 or 10) of the number of adaptive parameters number of parameters in the model [74, 75]. Even when we consider all 15,096 the samples in AGP as of May 2017 which mostly are collected from feces, the number of training samples is still less than 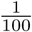 of the total number of trainable parameters, *m*. As an alternative, we instead propose to train a read level model, `Read2Pheno` which can leverage millions of reads as training examples from only dozens of samples, and then aggregate the learned information for a sample level prediction. One of the limitations of our model is that the model is not optimized for sample level prediction. However, through our experiments described here, we show that our proposed training strategy can pick up informative pattern to find 16S rRNA read and phenotype association as well as highlight informative regions.

In addition, unlike conventional OTU based methods, no preprocessing steps such as alignment/OTU grouping are required. Furthermore, intermediate layers outputs can be used for ordination and highlight informative regions in the sequences. Although Oligotyping can be used to resolve closely related 16S rRNA reads and explore the informative positions and label association, human supervision and alignment process are required to identify the cutoff entropy threshold in closely related reads of interest. Our model can be considered as an end-to-end model which takes raw 16S rRNA reads as input, learns informative genetic content and label association and outputs classification results. The learned knowledge can be extracted by intermediate layer outputs especially the attention weights. And the model is robust to deletions and insertions (as shown in Figure 5, our attention model can implicitly identify relevant genetic content across different unaligned sequences from the same genus). This work revealed the potential of deep learning models in phenotype prediction and interpretation. In the future, when a deep neural network model is better tuned for sample level prediction and when more training samples are made available, the method has the potential to outperform the existing OTU/ASV based methods.

To further show how the attention weights based model interpretation compares to other related method, We run the Oligotyping [41] software package on our Prevotella visualization set used in Section *Prevotella* Case Study. *Oligotyping* uses Shannon entropy to analyze closely related 16S rRNA sequences and find mutations that best explain sample variables (e.g. phenotypes). However, *Oligotyping* needs more human interaction and supervision for this task. For example, a user needs to determine how many oligotypes to find. Appendix Oligotypes for *Prevotella* reads shows the oligotypes (highlighted in black) found by their software for our American Gut derived Prevotella example. As shown in Appendix Oligotypes for *Prevotella* reads, *Oligotyping* for 7 positions picks 13, 54, 65, 83, 91, 96, and 98. In our model, shown in Figure 3, nucleotide positions 13, 54, 91, 96, 98 within the 16s rRNA V4 region are in highlighted region, which represent greater attention. Notably, position 13 is in a dim but still highlighted regions, 65 and 83 are near highlighted regions, and in fact, 83 is near two more. Thus, there is an apparent relationship between the attention that the neural network learns and the highest entropy positions that are learned. For each body site, there are multiple oligotypes per body site; therefore, it is hard to discern which high entropy position is most important to identifying which body site.

Several factors contribute to the discrepancy between our model’s high attention nucleotide positions and the highest entropy positions used for oligotyping. First, we smooth the attention weights with a 9-mer moving average. As a result, the attention weights are a regional approximation, i.e. not at precise positions, and the weights taper off at the ends of the read, due to edge convolutional effects. Second, *Oligotyping* only calculates the entropy of each position in Prevotella reads, while the attention weights learn the weighting of attention of positions that are *specifically* important to the classification task—in this case, body site prediction. A practical drawback of the *Oligotyping* approach is that, a user must plot the distribution of different oligotypes for each phenotype to see if there is a common oligotype for that phenotype, e.g., as in Appendix Oligotypes and body site association for *Prevotella*. From Appendix Oligotypes and body site association for *Prevotella*, the gut (fecal) oligotypes are evidently distinct from the other body sites, while the oral and skin oligotype distributions are relatively similar. Therefore, gut vs. other sites could be distinguished with oligotypes but more nucleotides would be needed to discern oral/skin. By contrast, the attention model does not require manual adjustment to find important regions. As the neural network classifies different phenotypes, the network learns the regions that are most important for this task. Attention modeling is thus the converse of oligotyping: attention weights recognize the informative reads/regions through body site classification and therefore can highlight the regions that distinguish body sites. Post-processing is required in the user’s end to extract the attention weights of sequences of interest for visualization. Conversely, preprocessing is required for *Oligotyping* users to identify reads of interest from a certain taxon and align them to learn oligotypes. Moreover, the attention weights for any combination of phenotype or taxa-phenotype can be visualized (see Figure 3), and it is immediately clear from the visualization what are the most informative sequence regions—as well as their relative nucleotide variability—for the classification being learned. Notably, with the attention model, the highlighted attention regions are slightly offset or in between high-entropy positions. Regions at or near high-density high-entropy positions (i.e., regions that have many high-entropy positions) are weighted with higher attention. We can see that highlighted attention maps to 4 sequence regions: nucleotide position ranges 12-18, 53-62, 74-81, and 90-98. Whereas region 4 (nt 90-98) helps identify oral reads and skin reads as opposed to gut, regions 1, 2, and 3 are helpful in distinguishing gut samples from other samples. Each body site may be identified by a combination of regions. For example, regions 2, 3, and 4 are useful to identify that a sample was taken from skin, while regions 1 and 4 are most useful for identifying samples from the oral cavity, and 1 and 3 are most useful for identifying gut samples.

Accordingly, the efficiency of attention model interpretability contrasts favorably compared to *Oligotyping* studies. As noted above, with *Oligotyping*, one must choose the number of nucleotides to examine. We chose 7 nucleotides, for example, since those represented nucleotide positions from all 4 regions in which we found attention. Moreover, there is no guarantee that *Oligotyping* will be able to succeed at all desired classification/discrimination tasks. For example, as Appendix Oligotypes and body site association for *Prevotella* shows, while the gut has a more distinct pattern of oligotypes as opposed to oral and skin samples, oral and skin samples show very little difference in oligotype patterns. As such, it is practically difficult to perform the classification task using only learned oligotypes. However, we do know, and have been able to quantify and show, the accuracy achieved by the attention model and its associated region discovery.

In addition to training and evaluating attention-based models on phenotype data, we have further considered a comprehensive data set for taxonomic classification, the SILVA rRNA database. SILVA data allowed us to evaluate how our attention model can be used to explore the structure of full length 16S sequences by training the model for a taxonomic classification task. (We can think of this as as `read2taxa`, but we are still using the same `Read2Pheno` model that we have developed.). Our model achieves comparable performance to the superior *k*-mer based method and pays attention to different regions for genus classification. This indicates that parts of the 16S rRNA gene, for a given 16S rRNA sequence from a genus, are important for distinguishing one genus from other genera. Notably, *Salmonella* has higher weights on V2 and V4. This can inform future 16S rRNA study designs. Because the V2 region is informative (i.e., has higher attention weights), investigators should design and use primers to target the V2 region to augment more commonly used V3/V4 primers. Some genera such as *Pirellula* have attention within some regions (V2 and V4) but also have higher than normal attentions at other sites, showing that more regions can be used to discern this genus. These predictions are consistent with evidence in the experimental literature. In particular, previous studies have shown that V1-V3 region are better at distinguishing *Escherichia*/*Shigella* [76], and we show a high attentions within the V2 and V3 for this group. The model also predicts that *Methanobacterium, Thermococcus*, and other Archaea have attention weighting at V4-V5. And, indeed, the V4-V5 regions have already been shown to have superior recognition of *Archaea* [77]. Accordingly, the `Read2Pheno` model’s predictions can serve as a starting point for identifying which primers may be optimally used to target various genera.

Going forward, we intend to adapt this model to explore metagenomic data, going beyond amplicon sequencing of specific marker genes to include the whole genome or sub-genomic multiple-gene regions from organisms in microbiome samples. To do so will require overcoming challenges such as memory limitations and the potential inability of neural networks to capture long distance dependencies due to gradient vanishing [78]. We are thus exploring the use of smaller batch size or more efficient data structure and better optimization strategy need to be applied to train the model for full length bacteria sequences. We are working to also achieve superior accuracy through approaches such as self-supervised pretraining and transfer learning, which have proven to be successful in the NLP literature [79, 80].

## Conclusion

In conclusion, we propose an integrated deep learning model that exploits CNNs, RNNs, and attention mechanisms to perform read and sample-level environmental prediction `and` extract interesting features. We show that such an alignment-free model can easily encode sequences/reads into dense and meaningful representations, and it can extract important sequence features while being robust to insertions and deletions. The `Read2Pheno` model we develop herein thus provides an exploratory way to understand microbial data which is helpful for future microbial study. In particular, we have shown in computational experiments that the deep, attention-based neural network `Read2Pheno` model can accomplish diverse tasks, learning informative nucleotides to (i) predict the body sites from which human microbiomic samples are extracted, (ii) discriminate between samples from individuals with positive and negative diagnoses for IBD, and (iii) identify the taxonomic labels of whole sequence 16s rRNA data. Moreover, we have shown that our attention-based DNN can not only predict, but can also be readily interpreted to obtain further insight. In addition, we have shown that we can interpret the intermediate outputs in the neural network model and generate visualization of the read embedding vectors, and, thereby discover important taxa and regions of sequences which are highly associated with the sample phenotype/taxonomic label. We show that our deep learning model can be used to explore the 16S rRNA nucleotide structure and its association with phenotype and taxonomy by learning the high-level features from *k*-mers and their sequential order. Our paper further provides an alternative way to train deep neural networks when the number of samples are relatively small, by training a read-level classifier instead of a sample-level classifier—while still producing comparable classification results. And, even in cases where our proposed attention-based neural network modeling framework fails to provide superior classification performance, we demonstrate that the results are nonetheless easy to interpret, such that we may extract important biologically and clinically relevant information from complex sequence data.

## Acknowledgments

We would like to thank Google LLC and Burwood Group, Inc. for their support. Google LLC provided us their Google Cloud Platform to train and validate our deep learning model. Burwood Group, Inc partnered with Google provide technical support and system configuration on Google Cloud Platform. In addition, work reported here was partially run on hardware supported by Drexel’s University Research Computing Facility (URCF). This work used the Extreme Science and Engineering Discovery Environment (XSEDE) [55], which is supported by National Science Foundation grant number ACI-1548562. Specifically, it used the Bridges system, which is supported by NSF award number ACI-1445606, at the Pittsburgh Supercomputing Center (PSC).

## A Appendix

### A.1 One-hot coding map

The map is adapted from Nomenclature for Incompletely Specified Bases in Nucleic Acid Sequences (http://www.sbcs.qmul.ac.uk/iubmb/misc/naseq.html)

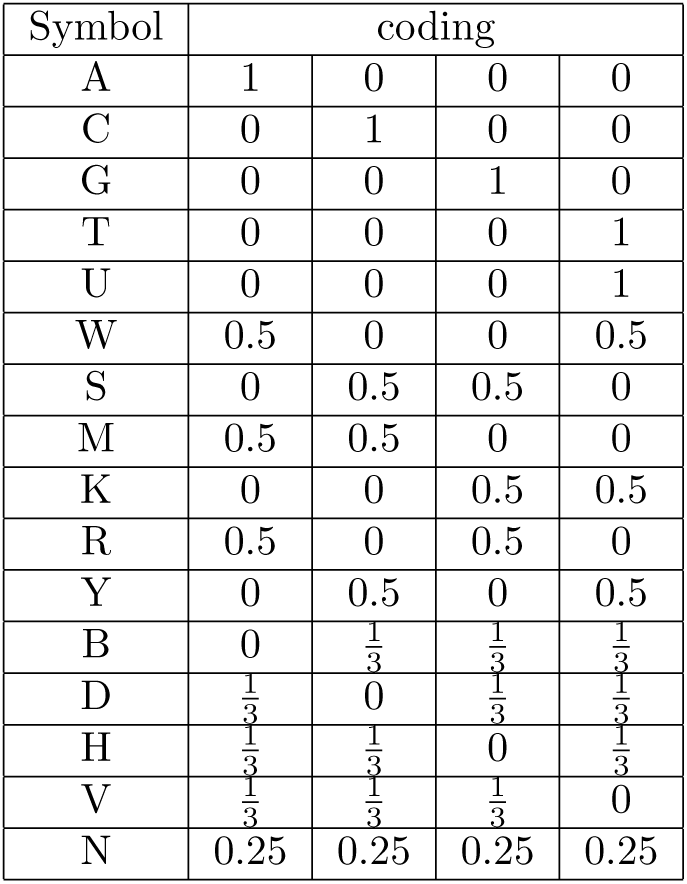

| Symbol | coding |  |  |  |
| --- | --- | --- | --- | --- |
| A | 1 | 0 | 0 | 0 |
| C | 0 | 1 | 0 | 0 |
| G | 0 | 0 | 1 | 0 |
| T | 0 | 0 | 0 | 1 |
| U | 0 | 0 | 0 | 1 |
| W | 0.5 | 0 | 0 | 0.5 |
| S | 0 | 0.5 | 0.5 | 0 |
| M | 0.5 | 0.5 | 0 | 0 |
| K | 0 | 0 | 0.5 | 0.5 |
| R | 0.5 | 0 | 0.5 | 0 |
| Y | 0 | 0.5 | 0 | 0.5 |
| B | 0 | $\frac{1}{3}$ | $\frac{1}{3}$ | $\frac{1}{3}$ |
| D | $\frac{1}{3}$ | 0 | $\frac{1}{3}$ | $\frac{1}{3}$ |
| H | $\frac{1}{3}$ | $\frac{1}{3}$ | 0 | $\frac{1}{3}$ |
| V | $\frac{1}{3}$ | $\frac{1}{3}$ | $\frac{1}{3}$ | 0 |
| N | 0.25 | 0.25 | 0.25 | 0.25 |

### A.2 Sample-level predictor: Majority vote method

The read caller threshold is 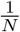 where N is the number of classes.

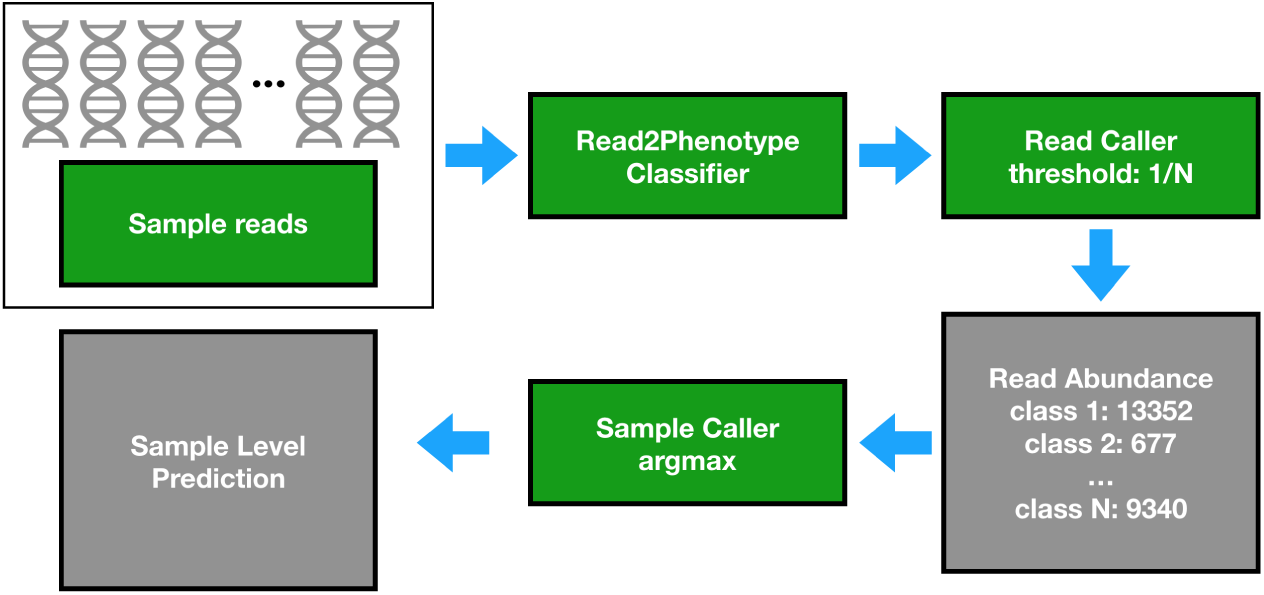

### A.3 Sample-level predictor: Sample-level embedding

Sample level embedding is calculated by averaging all the read level embedding vectors per sample. Then a random forest classifier is trained based on the sample embedding matrix (a *N* by *N*_*h*_ matrix where *N* is the total number of samples in training set and *N*_*h*_ is the number of hidden nodes in Bi-LSTM layer). Once the sample level random forest classifier is trained, this model can be used to perform sample level classification by taking the sample embedding vectors as input. The training and testing process are labeled by red and blue arrows correspondingly.

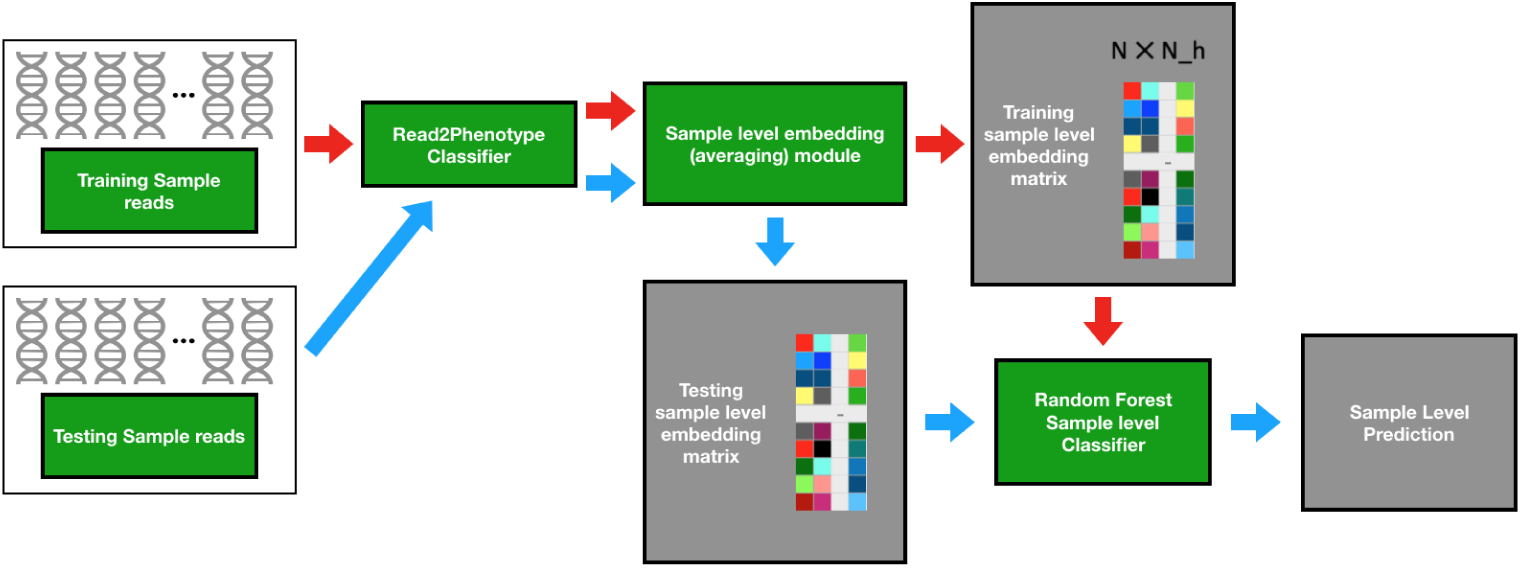

### A.4 Sample-level predictor: Pseudo-OTU

Training reads are clustered into *N*_*clusters*_ = 1000 clusters (“Pseudo OTUs”) by `Read level embedding clustering module` using a *k* −*means* algorithm. Then all training reads per sample are mapped to the closest “Pseudo OTUs” to form “Pseudo OTUs” abundance table. Similar to sample-level embedding method, a random forest classifier can be trained to perform sample level prediction using such “Pseudo OTUs” table (The training and testing process are labeled by red and blue arrows correspondingly).

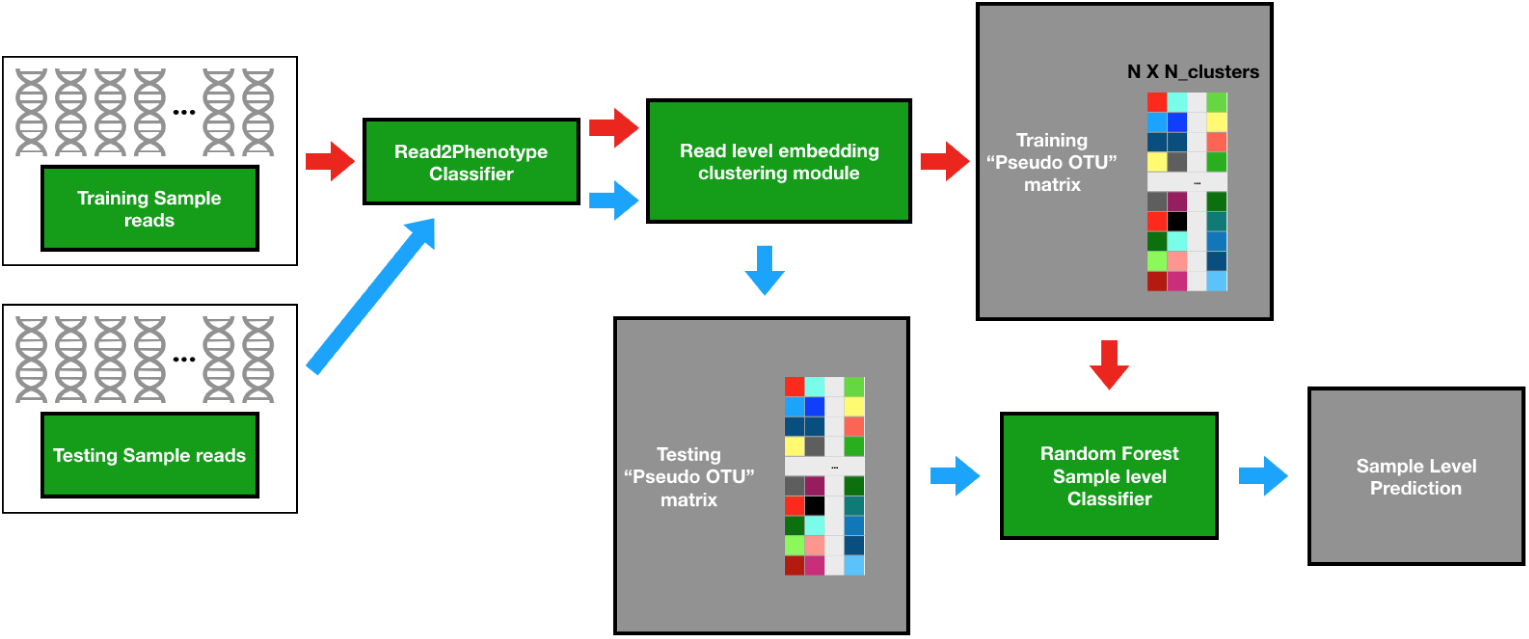

### A.5 Overall training and testing experiment

Samples are split into train and test set. Training set is used to train a `Read2Pheno` classifier and a sample-level predictor. The testing set is used to evaluate the performance.

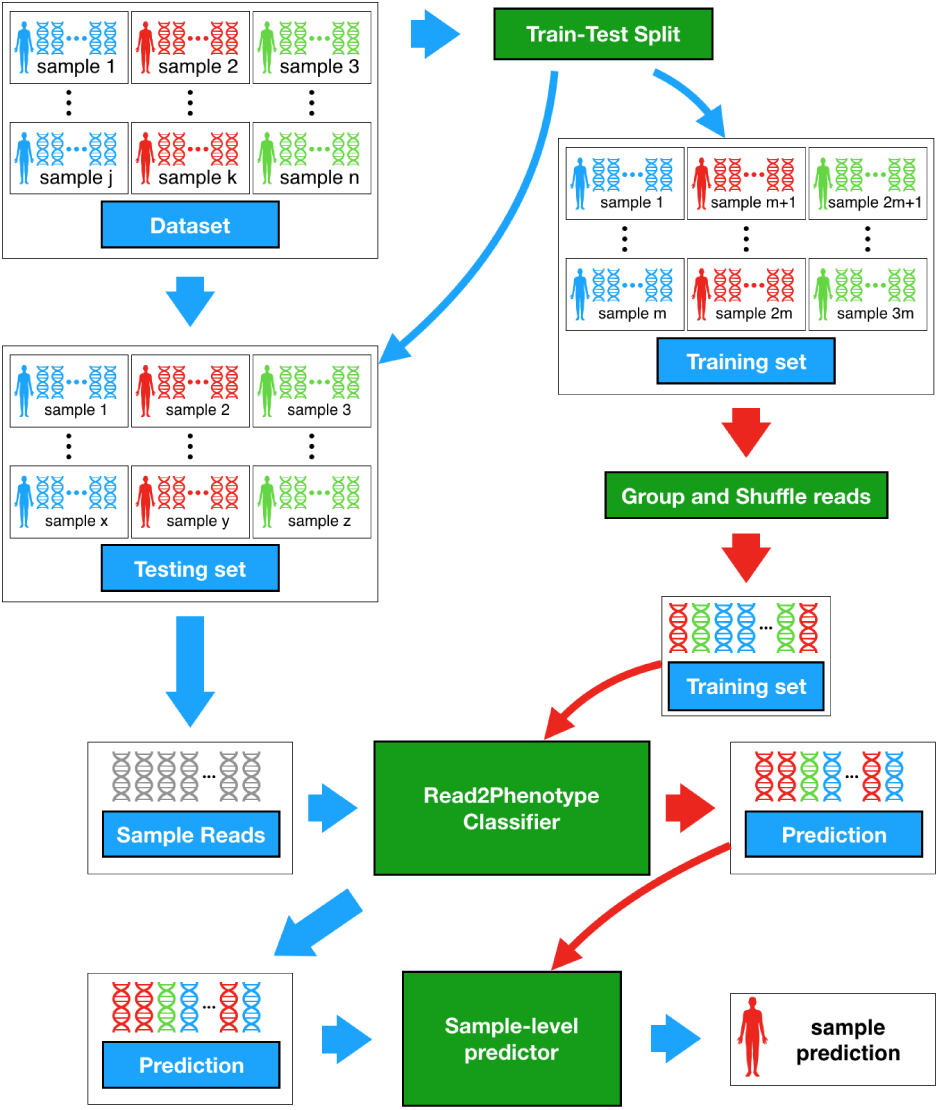

### A.6 Read2Pheno training process

All reads in the training set are labeled by the sample level label (the body site the original sample was collected from). Then the reads are grouped together and shuffled for training the `Read2Pheno` classifier.

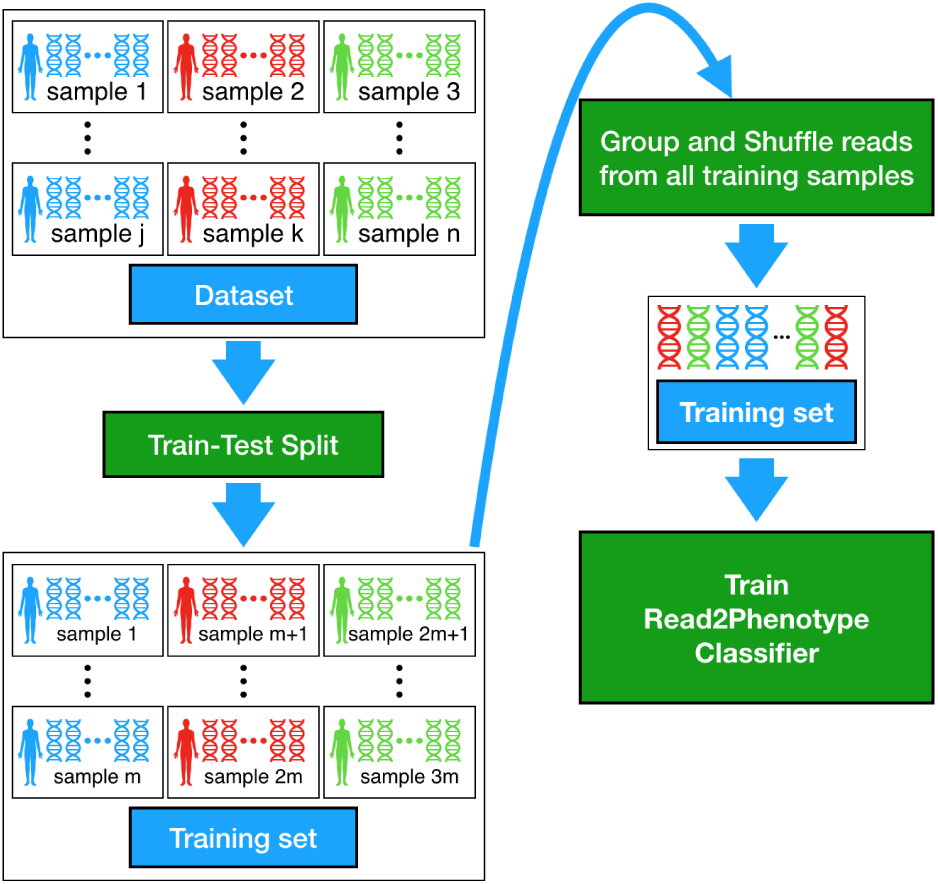

### A.7 Hyperparameter search space table

The best set of parameters is *{*number of conv filters, *N*_*c*_: 256, number of units in LSTM *N*_*h*_: 64, dropout probability for Dropout Layer: 0, learning rate: 0.001} for read level prediction on 5-fold cross validation of training data. The window size of convolutional layers, *W*, is set to 9 and the number of hidden nodes in attention layer, *N*_*a*_, is set to 16 in the default setting.

| Hyperparameter | Value |
| --- | --- |
| number of conv filters, $N_c$ | 128, 256 |
| number of units in LSTM, $N_h$ | 64, 128 |
| dropout probability for Dropout Layer | 0, 0.25 |
| learning rate | 0.001, 0.005 |

### A.8 Training data size effect of Read2Pheno classifier

Sample-level testing accuracy for different training data size of `Read2Pheno` classifier.

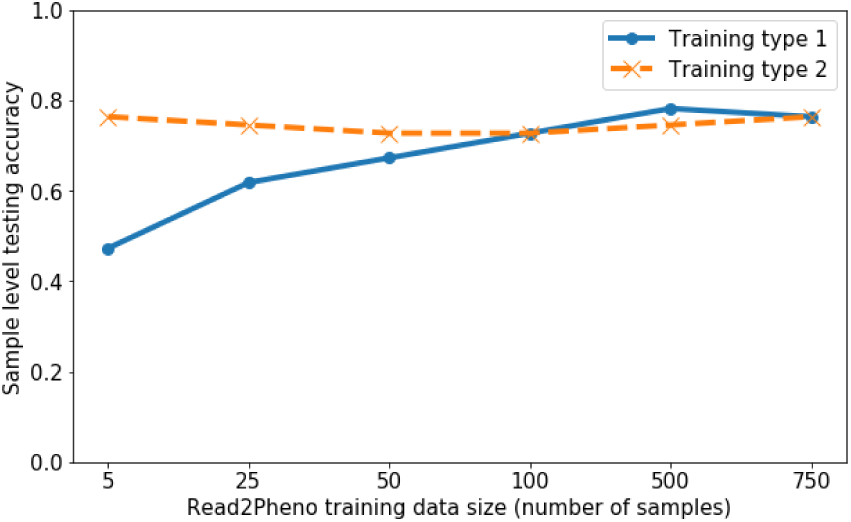

### A.9 2D Visualization of *Prevotella* reads

2-D projection of embedded *Prevotella* read vectors. Red markers represent fecal reads, green markers represent oral reads and black markers represent skin reads. The color of a ‘×’ represents the predicted body site. If the predicted body site is the same as the true body site, then ‘×’ s are not visible. The body site prediction accuracy for *Prevotella* visualization reads is 0.9131

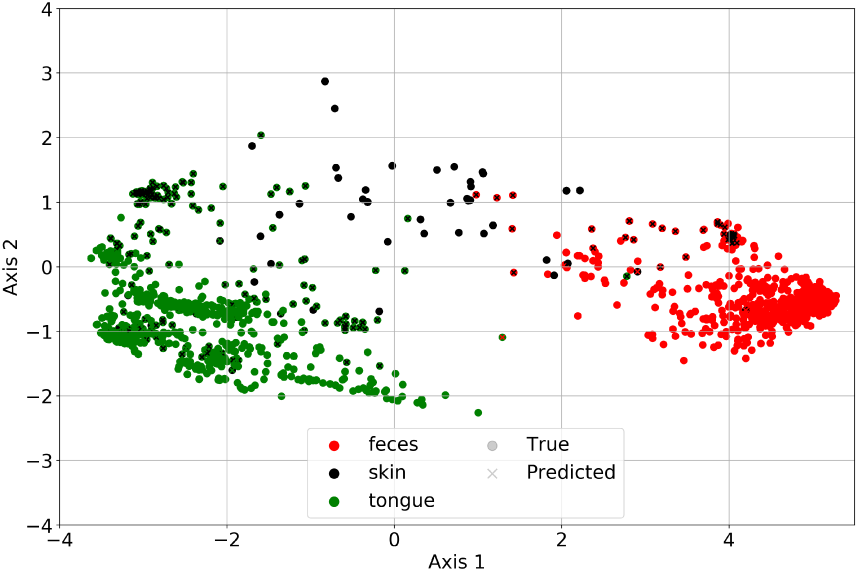

### A.10 Oligotypes for *Prevotella* reads

Top 7 Oligotypes found in *Prevotella* reads. The number on the left hand side of the figure shows the number of reads having a certain type of *Oligotyping* patterns (the nucleotide combination in black positions)

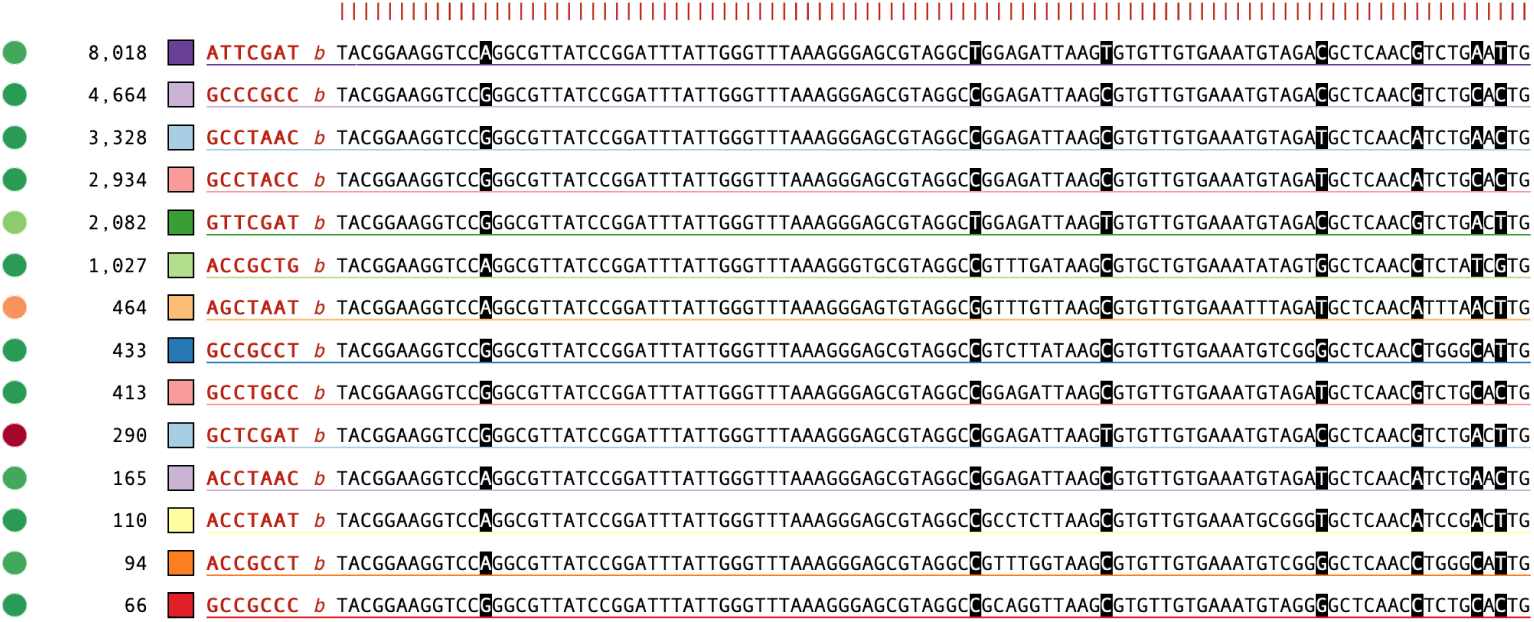

### A.11 Oligotypes and body site association for *Prevotella*

Oligotypes and body site association [41] of *Prevotella* reads. Different 7-oligotypes configurations can be found in different body sites. Fecal samples have a distinctive configuration vs. oral and skin samples.

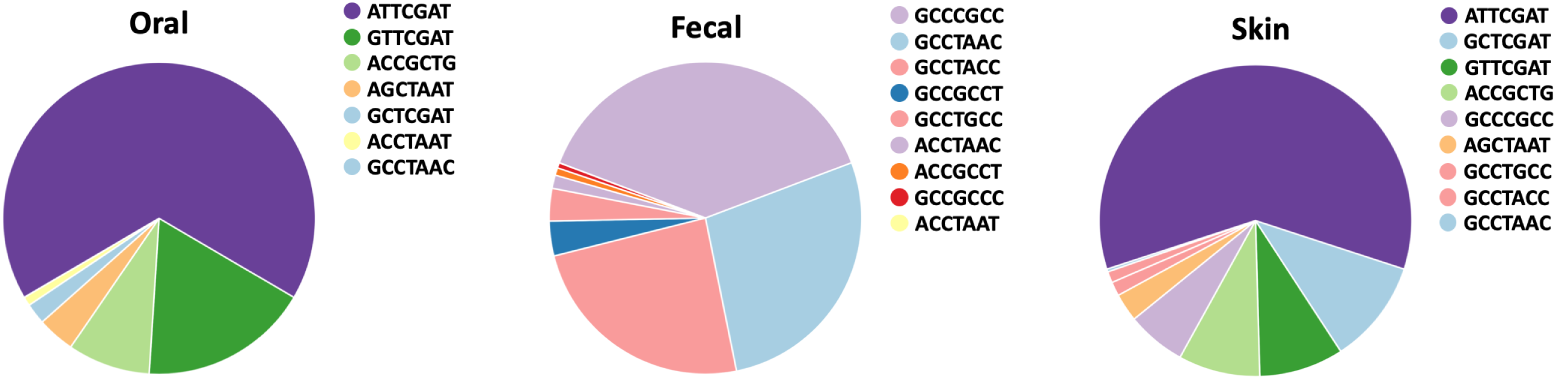

### A.12 2-D projection of embedded read vectors for Gevers dataset

2-D projection of embedded read vectors from all visualization samples (A), disease-positive samples (B), and disease-negative samples. ‘Δ’ s are reads from the control (health) samples and ‘○’ s are reads from the Crohn’s disease samples. The neural network is learning the 16S rRNA gene association to taxonomy and phenotype without the access to taxonomy label of the reads.

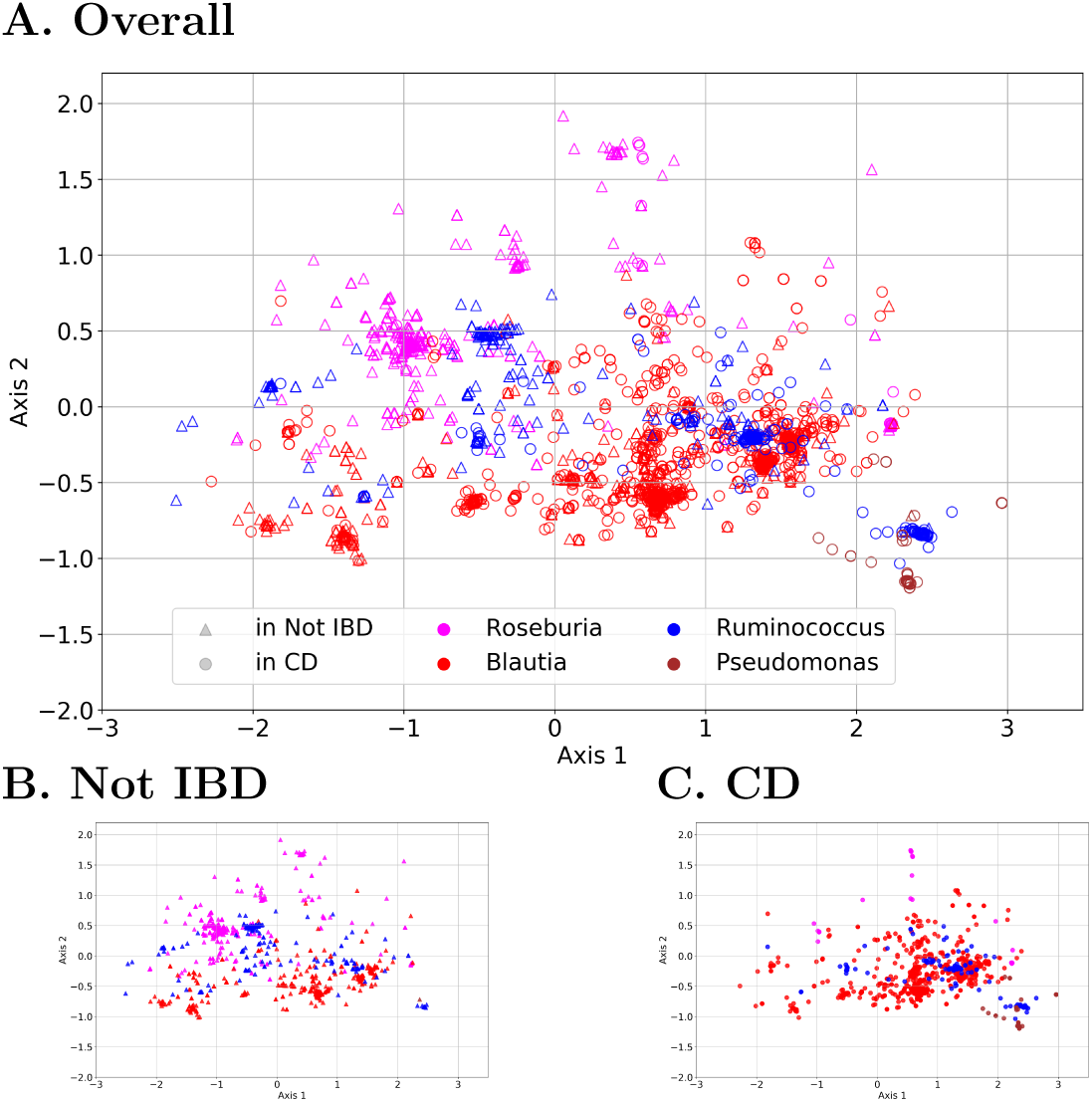

### A.13 Case studies of Gevers dataset

To understand which features facilitate class separation, we inspect the read embedded vectors for *Ruminococcus* and *Blautia* that separated well for phenotype. We also demonstrate interpretability of the attention mechanism by inspecting their attention weights.

### A.14 Average attention weights of *Enterobacter* testing sequences

Top: Average attention weights for all strains of *Enterobacter* without alignment; Bottom: Attention weights with alignment. This figure shows that our model can implicitly perform alignment to the sequence so that the attention was paid to similar position after alignment.

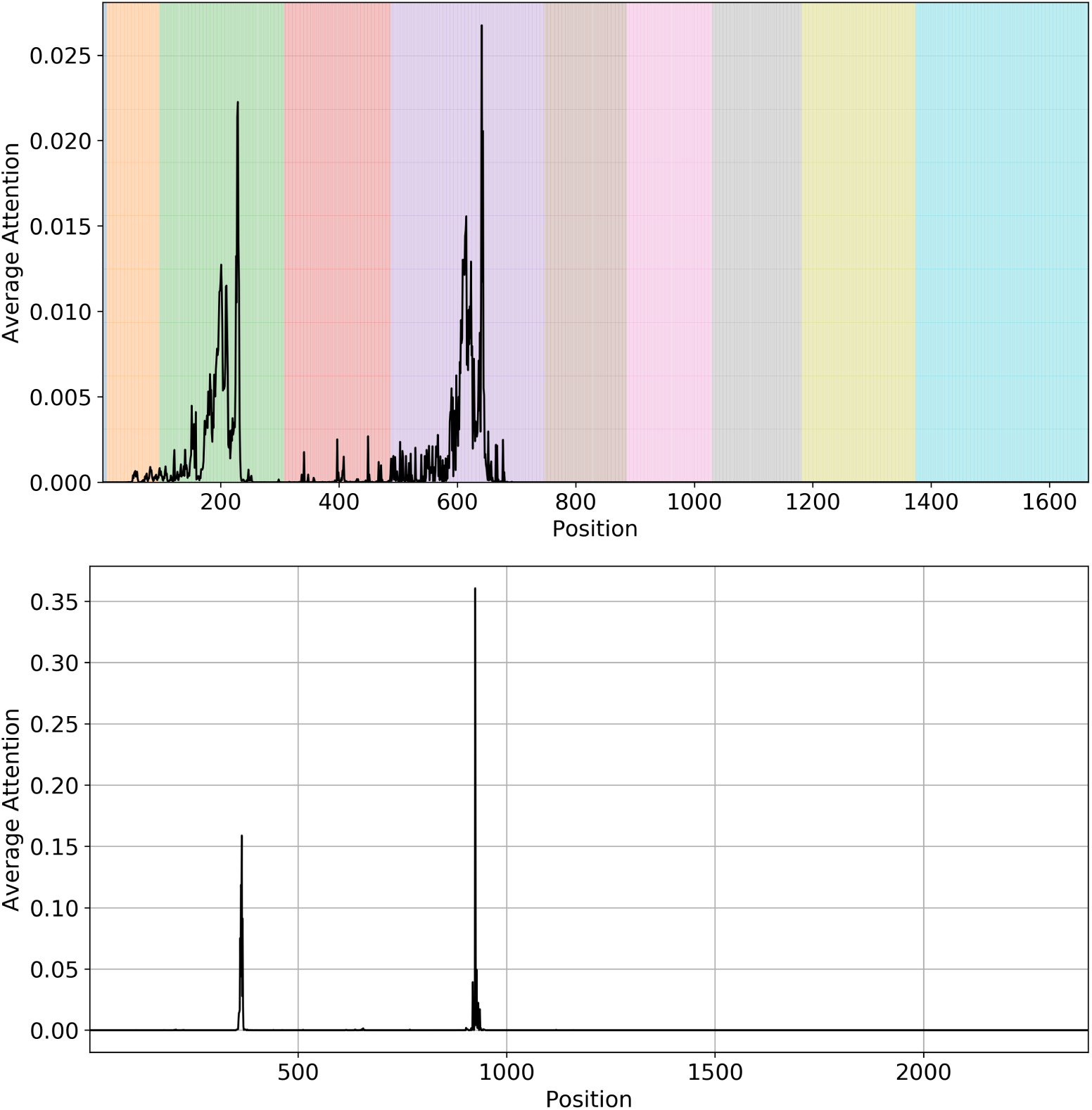

### A.15 Secondary structure and attention weights of a *Pseudomonas aeruginosa* sequence

The attention weights (top) on a real *Pseudomonas aeruginosa* sequence provided by [64]. The positions that have an attention weight greater than the mean attention weight cross the whole sequence is highlighted on the secondary structure figure (bottom).

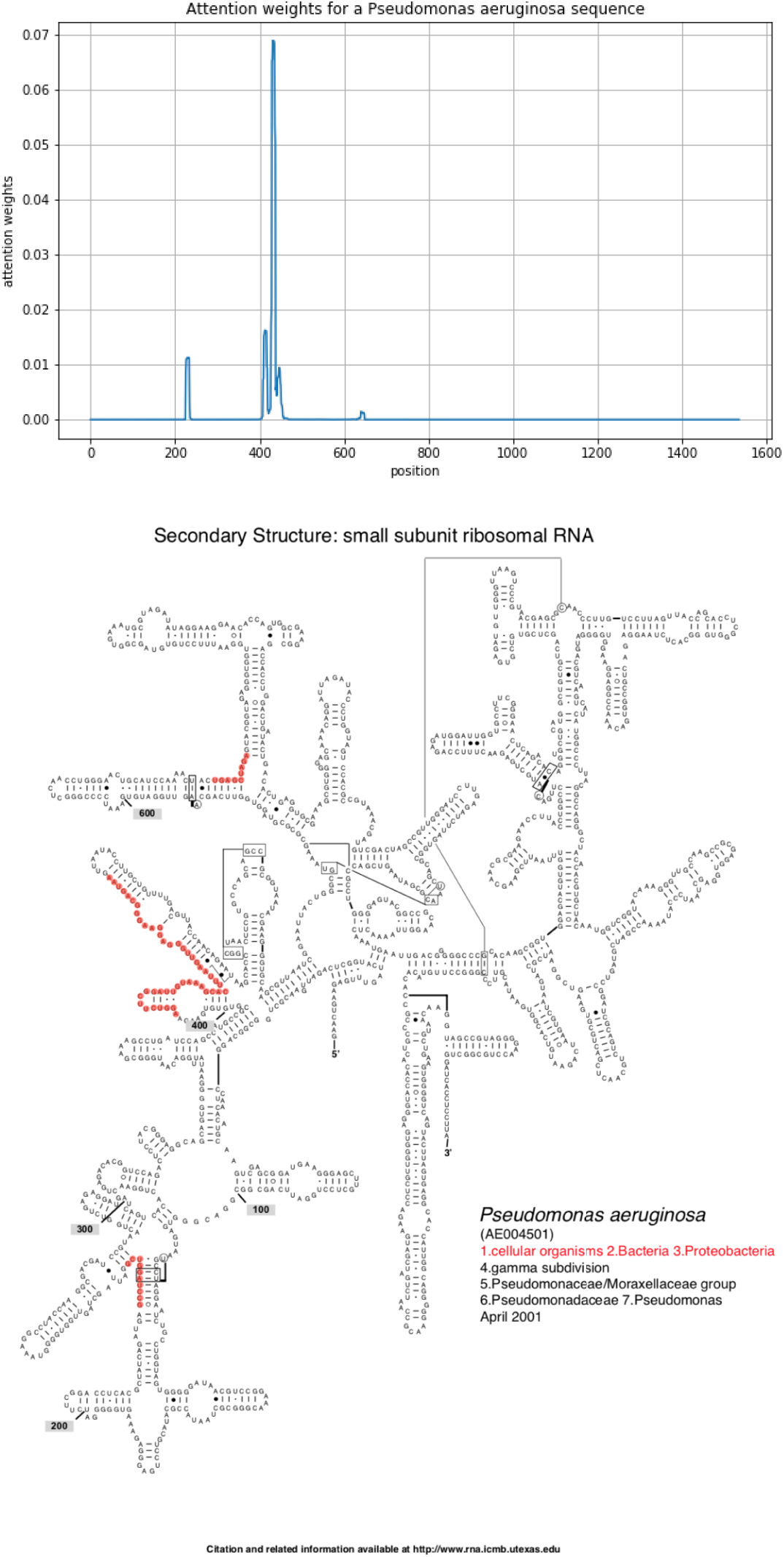

### A.16 Mean attention weights and multiple sequence alignment results aligned for *Pseudomonas* and *Enterobacter*

### A.17 Attention weights on variable regions per genus

## Footnotes

1 A python implementation of the proposed model is available at https://github.com/EESI/sequence_attention.

## References

1. Jose A. Navas-Molina, Embriette R. Hyde, Jon G. Sanders, and Rob Knight. The microbiome and big data. Current Opinion in Systems Biology, 4:92 – 96, 2017. Big data acquisition and analysis • Pharmacology and drug discovery.

2. A. Bernhard, D. Colbert, James McManus, and K. Field. Microbial community dynamics based on 16s rrna gene profiles in a pacific northwest estuary and its tributaries. FEMS microbiology ecology, 52 1:115–28, 2005.

3. Cindy H. Nakatsu, Muruleedhara N. Byappanahalli, and Meredith B Nevers. Bacterial community 16s rrna gene sequencing characterizes riverine microbial impact on lake michigan. Frontiers in Microbiology, 10, 2019.

4. Eri Nishiyama, Koichi Higashi, H. Mori, K. Suda, H. Nakamura, S. Omori, S. Maruyama, Y. Hongoh, and K. Kurokawa. The relationship between microbial community structures and environmental parameters revealed by metagenomic analysis of hot spring water in the kirishima area, japan. Frontiers in Bioengineering and Biotechnology, 6, 2018.

5. Renato Pedron, A. Esposito, Irene Bianconi, E. Pasolli, Adrian Tett, F. Asnicar, M. Cristofolini, N. Segata, and O. Jousson. Genomic and metagenomic insights into the microbial community of a thermal spring. Microbiome, 7, 2019.

6. Elizabeth M. Ross, Peter J. Moate, Leah C. Marett, Ben G. Cocks, and Ben J. Hayes. Metagenomic predictions: From microbiome to complex health and environmental phenotypes in humans and cattle. PLOS ONE, 8(9):1–8, 09 2013.

7. Dirk Gevers, Subra Kugathasan, Lee A Denson, Yoshiki Vázquez-Baeza, Will Van Treuren, Boyu Ren, Emma Schwager, Dan Knights, Se Jin Song, Moran Yassour, Xochitl C. Morgan, Aleksandar D. Kostic, Chengwei Luo, Antonio González, Daniel McDonald, Yael Haberman, Thomas D Walters, Susan S Baker, Joel R. Rosh, Michael C Stephens, Melvin B Heyman, James F. Markowitz, Robert N. Baldassano, Anne Marie Griffiths, Francisco Augusto Sylvester, David R. Mack, Sandra C. Kim, Wallace V. Crandall, Jeffrey S Hyams, Curtis Huttenhower, Rob Knight, and Ramnik J. Xavier. The treatment-näive microbiome in new-onset crohn’s disease. Cell host & microbe, 15 3:382–392, 2014.

8. Anna Paola Carrieri, Niina Haiminen, and Laxmi Parida. Host phenotype prediction from differentially abundant microbes using rodeo. In CIBB, 2016.

9. Ehsaneddin Asgari, Kiavash Garakani, Alice C McHardy, and Mohammad R K Mofrad. Micropheno: predicting environments and host phenotypes from 16s rrna gene sequencing using a k-mer based representation of shallow sub-samples. Bioinformatics, 34(13):i32–i42, 2018.

10. Guadalupe Navarro, Anukriti Sharma, Lara R. Dugas, Terrence Forrester, Jack A. Gilbert, and Brian T. Layden. Gut microbial features can predict host phenotype response to protein deficiency. Physiological Reports, 6(23):e13932. 2018.

11. Philip Hunter. Extended phenotype redux. how far can the reach of genes extend in manipulating the environment of an organism? EMBO reports, 10 3:212–5, 2009.

12. Chad M. Cullen, Kawalpreet K. Aneja, Sinem Beyhan, Clara E. Cho, Stephen Woloszynek, Matteo Convertino, Sophie J McCoy, Yanyan Zhang, Matthew Z Anderson, David Alvarez-Ponce, Ekaterina Smirnova, Lisa Karstens, Pieter C. Dorrestein, Hongzhe Li, Ananya Sen Gupta, Kevin K W Cheung, Jennifer Gloeckner Powers, Zhengqiao Zhao, and Gail L. Rosen. Emerging priorities for microbiome research. Frontiers in Microbiology, 11, 2020.

13. M. Fischbach. Microbiome: Focus on causation and mechanism. Cell, 174:785–790, 2018.

14. Tonya L Ward, Jake Larson, Jeremy Meulemans, Ben Hillmann, Joshua Lynch, D. Sidiropoulos, J. Spear, G. Caporaso, Ran Blekhman, R. Knight, R. Fink, and Dan Knights. Bugbase predicts organism-level microbiome phenotypes. bioRxiv, 2017.

15. K. Lu, Ridwan Mahbub, P. Cable, H. Ru, N. Parry, W. Bodnar, J. S. Wishnok, M. Stýblo, J. Swenberg, J. Fox, and S. Tannenbaum. Gut microbiome phenotypes driven by host genetics affect arsenic metabolism. Chemical Research in Toxicology, 27:172 – 174, 2014.

16. M. Stanislawski, D. Dabelea, Leslie A. Lange, B. Wagner, and C. Lozupone. Gut microbiota phenotypes of obesity. NPJ Biofilms and Microbiomes, 5, 2019.

17. J. B. Lynch and E. Y. Hsiao. Microbiomes as sources of emergent host phenotypes. Science, 365:1405 – 1409, 2019.

18. E. Ross, P. Moate, L. Marett, B. Cocks, and B. Hayes. Metagenomic predictions: From microbiome to complex health and environmental phenotypes in humans and cattle. PLoS ONE, 8, 2013.

19. A. Bhattacharjee, Dušan Veličković, T. Wietsma, Sheryl L. Bell, J. Jansson, K. Hofmockel, and C. Anderton. Visualizing microbial community dynamics via a controllable soil environment. mSystems, 5, 2020.

20. Jolinda Pollock, Laura Glendinning, Trong Wisedchanwet, and Mick Watson. The madness of microbiome: Attempting to find consensus “best practice” for 16s microbiome studies. Applied and Environmental Microbiology, 84, 2018.

21. Alexander Statnikov, Mikael Henaff, Varun Narendra, Kranti Konganti, Zhiguo Li, Liying Yang, Zhiheng Pei, Martin J. Blaser, Constantin F. Aliferis, and Alexander V. Alekseyenko. A comprehensive evaluation of multicategory classification methods for microbiomic data. Microbiome, 1(1):11, Apr 2013.

22. Karen Simonyan, Andrea Vedaldi, and Andrew Zisserman. Deep inside convolutional networks: Visualising image classification models and saliency maps. CoRR, abs/1312.6034, 2013.

23. Avanti Shrikumar, Peyton Greenside, Anna Shcherbina, and Anshul Kundaje. Not Just a Black Box: Learning Important Features Through Propagating Activation Differences. ArXiv, may 2016.

24. Karen Simonyan, Andrea Vedaldi, and Andrew Zisserman. Deep Inside Convolutional Networks: Visualising Image Classification Models and Saliency Maps. ArXiv, pages 1–8, 2013.

25. Jason Yosinski, Jeff Clune, Anh Nguyen, Thomas Fuchs, and Hod Lipson. Understanding Neural Networks Through Deep Visualization. ArXiv, jun 2015.

26. Seonwoo Min, Byunghan Lee, and Sungroh Yoon. Deep learning in bioinformatics. Briefings in Bioinformatics, page bbw068, jul 2016.

27. G. Ditzler, R. Polikar, and G. Rosen. Multi-layer and recursive neural networks for metagenomic classification. IEEE Transactions on NanoBioscience, 14:608–616, 2015.

28. Jack Lanchantin, Ritambhara Singh, Zeming Lin, and Yanjun Qi. Deep Motif: Visualizing Genomic Sequence Classifications. ArXiv, may 2016.

29. Laura Deming, Sasha Targ, Nate Sauder, Diogo Almeida, and Chun Jimmie Ye. Genetic Architect: Discovering Genomic Structure with Learned Neural Architectures. ArXiv, may 2016.

30. Ryan Poplin, Dan Newburger, Jojo Dijamco, Nam Ngoc Nguyen, Dion Loy, Sam Gross, Cory Y McLean, and Mark A. DePristo. Creating a universal snp and small indel variant caller with deep neural networks. 2017.

31. Moritz Hess, Stefan Lenz, Tamara J. Blätte, Lars Bullinger, and Harald Binder. Partitioned learning of deep Boltzmann machines for SNP data. Bioinformatics, 33(20):3173–3180, oct 2017.

32. Akosua Busia, George E. Dahl, Clara Fannjiang, David H. Alexander, Elizabeth Dorfman, Ryan Poplin, Cory Y. McLean, Pi-Chuan Chang, and Mark DePristo. A deep learning approach to pattern recognition for short dna sequences. bioRxiv, 2018.

33. Daniel Quang and Xiaohui Xie. Danq: A hybrid convolutional and recurrent deep neural network for quantifying the function of dna sequences. 44:gkw226, 04 2016.

34. Yifei Chen, Yi Li, Rajiv Narayan, Aravind Subramanian, and Xiaohui Xie. Gene expression inference with deep learning. Bioinformatics, 32(12):1832–1839, jun 2016.

35. Chieh Lo and Radu Marculescu. Metann: accurate classification of host phenotypes from metagenomic data using neural networks. BMC Bioinformatics, 20, 2019.

36. Thanh Hai Nguyen, Edi Prifti, Yann Chevaleyre, Nataliya Sokolovska, and Jean-Daniel Zucker. Disease classification in metagenomics with 2d embeddings and deep learning. ArXiv, abs/1806.09046, 2018.

37. D. Reiman, A. Metwally, J. Sun, and Y. Dai. Popphy-cnn: A phylogenetic tree embedded architecture for convolutional neural networks to predict host phenotype from metagenomic data. IEEE Journal of Biomedical and Health Informatics, pages 1–1, 2020.

38. Stephen Woloszynek, Zhengqiao Zhao, Jian Chen, and Gail L. Rosen. 16s rrna sequence embeddings: Meaningful numeric feature representations of nucleotide sequences that are convenient for downstream analyses. PLOS Computational Biology, 15(2):1–25, 02 2019.

39. Nathan Lapierre, Chelsea Ju, Guangyu Zhou, and Wei Wang. Metapheno: A critical evaluation of deep learning and machine learning in metagenome-based disease prediction. Methods, 2019.

40. W. James Murdoch, Chandan Singh, Karl Kumbier, Reza Abbasi-Asl, and B. Yu. Definitions, methods, and applications in interpretable machine learning. Proceedings of the National Academy of Sciences, 116:22071 – 22080, 2019.

41. A. Murat Eren, Löis Maignien, Woo Jun Sul, Leslie G. Murphy, Sharon L. Grim, Hilary G. Morrison, and Mitchell L. Sogin. Oligotyping: differentiating between closely related microbial taxa using 16s rrna gene data. Methods in Ecology and Evolution, 4(12):1111–1119, 2013.

42. Erki Aun, Age Brauer, Veljo Kisand, Tanel Tenson, and Maido Remm. A k-mer-based method for the identification of phenotype-associated genomic biomarkers and predicting phenotypes of sequenced bacteria. PLOS Computational Biology, 14(10):1–17, 10 2018.

43. Babak Alipanahi, Andrew Delong, Matthew T. Weirauch, and Brendan J. Frey. Predicting the sequence specificities of dna- and rna-binding proteins by deep learning. Nature Biotechnology, 33:831–838, 2015.

44. Jack Lanchantin, Ritambhara Singh, Beilun Wang, and Yanjun Qi. Deep Motif Dashboard: Visualizing and Understanding Genomic Sequences Using Deep Neural Networks. ArXiv, aug 2016.

45. Dzmitry Bahdanau, Kyunghyun Cho, and Yoshua Bengio. Neural machine translation by jointly learning to align and translate. CoRR, abs/1409.0473, 2015.

46. Zichao Yang, Diyi Yang, Chris Dyer, Xiaodong He, Alexander J. Smola, et al. Hierarchical attention networks for document classification. In HLT-NAACL, 2016.

47. Peng Zhou, Wei Shi, Jun Tian, Zhenyu Qi, Bingchen Li, Hongwei Hao, and Bo Xu. Attention-based bidirectional long short-term memory networks for relation classification. In ACL, 2016.

48. Qiao Liu, Haibin Zhang, Yifu Zeng, Ziqi Huang, and Zufeng Wu. Content attention model for aspect based sentiment analysis. In Proceedings of the 2018 World Wide Web Conference, WWW’18, pages 1023–1032, Republic and Canton of Geneva, Switzerland, 2018. International World Wide Web Conferences Steering Committee.

49. Daniel McDonald, Embriette R. Hyde, Justine W. Debelius, James T. Morton, Antonio González, Gail Ackermann, Alexander A. Aksenov, Bahar Behsaz, Caitriona Brennan, Yingfeng Chen, Lindsay DeRight Goldasich, Pieter C. Dorrestein, Robert R. Dunn, Ashkaan K Fahimipour, James A. Gaffney, Jack A. Gilbert, Grant Gogul, Jessica L. Green, Philip Hugenholtz, Greg C. Humphrey, Curtis Huttenhower, Matthew A Jackson, Stefan Janssen, Dilip V. Jeste, Lingjing Jiang, Scott T. Kelley, Dan Knights, Tomasz Kosciólek, Joshua Ladau, Jeff Leach, Clarisse Marotz, Dmitry Meleshko, Alexey V. Melnik, Jessica L. Metcalf, Hosein Mohimani, Emmanuel Montassier, Jose A Navas-Molina, Tanya T Nguyen, Shyamal Das Peddada, Pavel Pevzner, Katherine S. Pollard, Gholamali Rahnavard, A. Robbins-Pianka, Naseer Sangwan, Joshua Shorenstein, Larry Smarr, Se Jin Song, Timothy David Spector, Austin D. Swafford, Varykina G Thackray, Luke R Thompson, Anupriya Tripathi, Yoshiki Vázquez-Baeza, Alison F. Vrbanac, Paul E Wischmeyer, Elaine Wolfe, Qiyun Zhu, and Rob Knight. American gut: an open platform for citizen science microbiome research. mSystems, 3, 2018.

50. Christian Quast, Elmar Pruesse, Pelin Yilmaz, Jan Gerken, Timmy Schweer, Pablo Yarza, Jörg Peplies, and Frank Oliver Glöckner. The SILVA ribosomal RNA gene database project: improved data processing and web-based tools. Nucleic Acids Research, 41(D1):D590–D596, 11 2012.

51. Pelin Yilmaz, Laura Wegener Parfrey, Pablo Yarza, Jan Gerken, Elmar Pruesse, Christian Quast, Timmy Schweer, Jörg Peplies, Wolfgang Ludwig, and Frank Oliver Glöckner. The SILVA and “All-species Living Tree Project (LTP)” taxonomic frameworks. Nucleic Acids Research, 42(D1):D643–D648, 11 2013.

52. Colin Raffel and Daniel P. W. Ellis. Feed-forward networks with attention can solve some long-term memory problems. ArXiv, abs/1512.08756, 2015.

53. J Gregory Caporaso, Justin Kuczynski, Jesse Stombaugh, Kyle Bittinger, Frederic D Bushman, Elizabeth K Costello, Noah Fierer, Antonio Gonzalez Peña, Julia Goodrich, Jeffrey I Gordon, Gavin Huttley, Scott T Kelley, Dan Knights, Jeremy E Koenig, Ruth Ley, Catherine Lozupone, Daniel Mcdonald, Brian D Muegge, Meg Pirrung, and Rob Knight. Qiime allows analysis of high-throughput community sequencing data. nat met 7: 335-336. Nature methods, 7:335–6, 04 2010.

54. James R. Cole, Qiong Wang, Jordan A. Fish, Benli Chai, Donna M. McGarrell, Yanni Sun, C. Titus Brown, Andrea Porras-Alfaro, Cheryl R. Kuske, and James M. Tiedje. Ribosomal Database Project: data and tools for high throughput rRNA analysis. Nucleic Acids Research, 42(D1):D633–D642, 11 2013.

55. J. Towns, T. Cockerill, M. Dahan, I. Foster, K. Gaither, A. Grimshaw, V. Hazlewood, S. Lathrop, D. Lifka, G. D. Peterson, R. Roskies, J. R. Scott, and N. Wilkins-Diehr. Xsede: Accelerating scientific discovery. Computing in Science Engineering, 16(5):62–74, Sep. 2014.

56. Gavin E Crooks, Gary Hon, John-Marc Chandonia, and Steven E Brenner. Weblogo: a sequence logo generator. Genome research, 14:1188–90, 07 2004.

57. ImportanceOfBeingErnest. sequence logos in matplotlib: aligning xticks, March 2017.

58. Benjamin J. Callahan, Paul J. McMurdie, Michael J. Rosen, Andrew W. Han, Amy Jo A Johnson, and Susan P. Holmes. Dada2: High resolution sample inference from illumina amplicon data. Nature methods, 13:581 – 583, 2016.

59. Josef Wagner, Anthony G. Catto-Smith, Donald J.S. Cameron, and Carl D. Kirkwood. Pseudomonas Infection in Children with Early-onset Crohn’s Disease: An Association with a Mutation Close to PSMG1. Inflammatory Bowel Diseases, 19(4):|pE58–E59, 05 2012.

60. Peter De Cruz, Lani Prideaux, Josef Wagner, Siew C. Ng, Chris McSweeney, Carl Kirkwood, Mark Morrison, and Michael A. Kamm. Characterization of the gastrointestinal microbiota in health and inflammatory bowel disease. Inflammatory Bowel Diseases, 18(2):372–390, 2012.

61. Josef Wagner, Kirsty Short, Anthony G. Catto-Smith, Don J. S. Cameron, Ruth F. Bishop, and Carl D. Kirkwood. Identification and characterisation of pseudomonas 16s ribosomal dna from ileal biopsies of children with crohn’s disease. PLOS ONE, 3(10):1–7, 10 2008.

62. Bo Yang, Yong Wang, and Pei-Yuan Qian. Sensitivity and correlation of hypervariable regions in 16s rrna genes in phylogenetic analysis. BMC Bioinformatics, 17, 12 2016.

63. M. A. Miller, W. Pfeiffer, and T. Schwartz. Creating the cipres science gateway for inference of large phylogenetic trees. In 2010 Gateway Computing Environments Workshop (GCE), pages 1–8, Nov 2010.

64. Jamie J. Cannone, Sankar Subramanian, Murray N. Schnare, James R. Collett, Lisa M. D’Souza, Yushi Du, Brian Feng, Nan Lin, Lakshmi V. Madabusi, Kirsten M. Müller, Nupur Pande, Zhidi Shang, Nan Yu, and Robin R. Gutell. The comparative rna web (crw) site: an online database of comparative sequence and structure information for ribosomal, intron, and other rnas. BMC Bioinformatics, 3(1):2, Jan 2002.

65. Hilde Vinje, Trygve Almøy, Kristian Liland, and Lars Snipen. A systematic search for discriminating sites in the 16s ribosomal rna gene. Microbial informatics and experimentation, 4:|p2, 01 2014.

66. Himel Mallick, Eric Franzosa, Lauren Mclver, Soumya Banerjee, Alexandra Sirota-Madi, Aleksandar Kostic, Clary Clish, Hera Vlamakis, Ramnik Xavier, and Curtis Huttenhower. Predictive metabolomic profiling of microbial communities using amplicon or metagenomic sequences. Nature Communications, 10:3136–3146, 07 2019.

67. Simon Graspeuntner, Nathalie Loeper, Sven Künzel, John Baines, and Jan Rupp. Selection of validated hypervariable regions is crucial in 16s-based microbiota studies of the female genital tract. Scientific Reports, 8, 06 2018.

68. Zigui Chen, Pak Chun Hui, Mamie Hui, Yun Kit Yeoh, Po Yee Wong, Martin C. W. Chan, Martin C. S. Wong, Siew C. Ng, Francis K. L. Chan, and Paul K. S. Chan. Impact of preservation method and 16s rrna hypervariable region on gut microbiota profiling. mSystems, 4(1), 2019.

69. Daniel McDonald, Amanda Birmingham, and Rob Knight. Context and the human microbiome. Microbiome, 3(1):52, Nov 2015.

70. Catherine Lozupone and Rob Knight. Unifrac: a new phylogenetic method for comparing microbial communities. Applied and Environmental Microbiology, 71(12):8228–8235, 2005.

71. Daniel McDonald, Zhenjiang Xu, Embriette R. Hyde, and Rob Knight. Ribosomal rna, the lens into life. Cold Spring Harbor Laboratory Press for the RNA Society, 2015.

72. Francis Ha and Hanan Khalil. Crohn’s disease: a clinical update. Therapeutic Advances in Gastroenterology, 8(6):352–359, 2015. PMID: 26557891.

73. Victoria Pascal, Marta Pozuelo, Natalia Borruel, Francesc Casellas, David Campos, Alba Santiago, Xavier Martinez, Encarna Varela, Guillaume Sarrabayrouse, Kathleen Machiels, Severine Vermeire, Harry Sokol, Francisco Guarner, and Chaysavanh Manichanh. A microbial signature for crohn’s disease. Gut, 66(5):813–822, 2017.

74. C. M. Bishop. Pattern recognition and machine learning (information science and statistics). 2006.

75. C. Zhang, S. Bengio, M. Hardt, B. Recht, and Oriol Vinyals. Understanding deep learning requires rethinking generalization. ArXiv, abs/1611.03530, 2017.

76. Jethro S. Johnson, Daniel J Spakowicz, Bo young Hong, Lauren M. Petersen, Patrick Demkowicz, Lei Chen, Shana R. Leopold, Blake M. Hanson, Hanako O. Agresta, Mark B. Gerstein, Erica Sodergren, and George M. Weinstock. Evaluation of 16s rrna gene sequencing for species and strain-level microbiome analysis. Nature Communications, 10, 2019.

77. Ciara Willis, Dhwani K. Desai, and Julie Laroche. Influence of 16s rrna variable region on perceived diversity of marine microbial communities of the northern north atlantic. FEMS Microbiology Letters, 366, 2019.

78. D. Arpit, Bhargav Kanuparthi, Giancarlo Kerg, Nan Rosemary Ke, Ioannis Mitliagkas, and Yoshua Bengio. h-detach: Modifying the lstm gradient towards better optimization. ArXiv, abs/1810.03023, 2019.

79. J. Devlin, Ming-Wei Chang, Kenton Lee, and Kristina Toutanova. Bert: Pre-training of deep bidirectional transformers for language understanding. In NAACL-HLT, 2019.

80. A. Radford. Improving language understanding by generative pre-training. 2018.

