## Supplementary material for "Learning, Visualizing and Exploring 16S rRNA Structure Using an Attention-based Deep Neural Network": A13, A16 and A17: A13.pdf

### Case studies of Gevers dataset

To understand which features facilitate class separation, we again inspect the read embedded vectors for genera that separated well for phenotype. Notice in Figure 1, reads form two reasonably separated clusters, one for “CD” and “Not IBD” in both ***Ruminococcus*** (Left) and ***Blautia*** (Right) panels. Therefore, we further use ***Ruminococcus*** and ***Blautia*** as exemplary demonstrations of the interpretability of the attention mechanism.

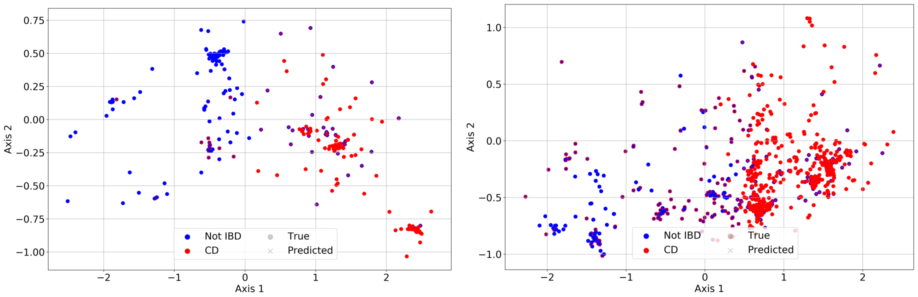


**Fig 1.** 2-D projection of embedded read vectors from ***Ruminococcus*** (Left) and ***Blautia*** (Right). Red markers represent “CD” reads, blue markers represent “Not IBD” reads. The color of a ‘×’ represents the predicted phenotype. If the predicted phenotype is the same as the true phenotype, then ‘×’s are not visible. The phenotype prediction accuracy for ***Ruminococcus*** and ***Blautia*** visualization reads are 0.84 and 0.74 respecitively

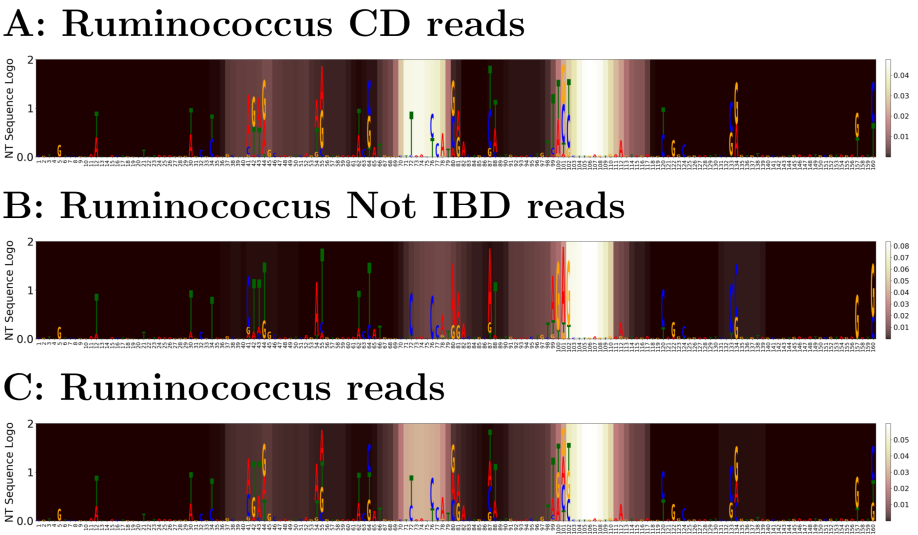


**Fig 2.** Comparison between average ***Ruminococcus*** reads attention and nucleotide frequency entropy in form of nucleotide sequence logo. A: “CD” reads; B: “Not IBD” reads; C: overall attention. In each phenotype, nucleotide frequency are scaled by the overall entropy for all ***Ruminococcus*** testing reads and plotted as sequence logo and averaged attention weights are represented by the color map (brighter colors represent larger attention weights)

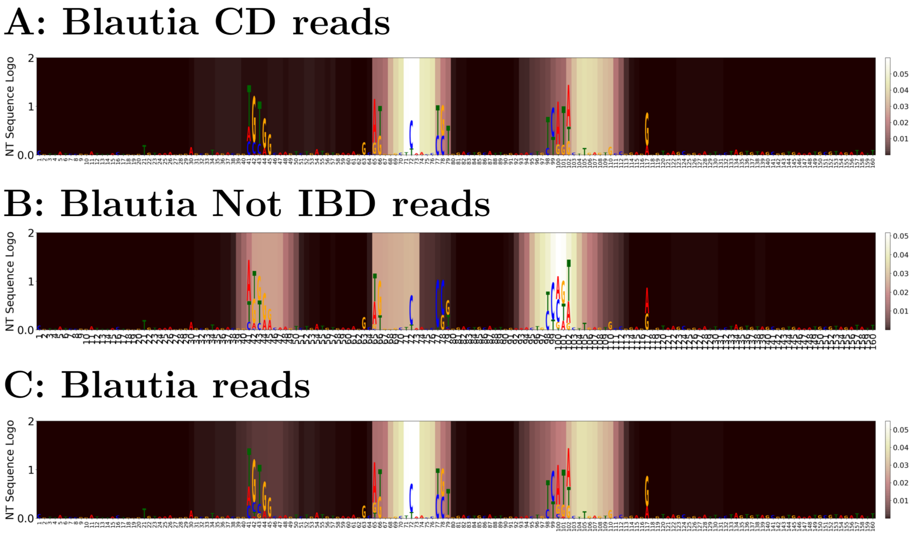


**Fig 3.** Comparison between average ***Blautia*** reads attention and nucleotide frequency entropy in form of nucleotide sequence logo. A: "CD" reads; B: "Not IBD" reads; C: overall attention. In each phenotype, nucleotide frequency are scaled by the overall entropy for all ***Blautia*** testing reads and plotted as sequence logo and averaged attention weights are represented by the color map (brighter colors represent larger attention weights)

To visualize the regions that are most informative to this classification, we inspect the input's attention weights as well as the nucleotide frequency. Figure 2 and 3 show the correlation between high entropy positions and high attention positions using the method described in "Model Interpretation and Read Visualization" section in the manuscript. For better visualization, the attention weights are smoothed by a moving average of window size of 9 (the size used in the convolutional filter in our deep learning model).

As shown in the figures, the model is more likely to pay attention to nucleotide variable regions that are informative. Te high attention weight region and the high nucleotide variance region coincides well with each other in Figure 3. However, there is some shift between the high attention weight region and the high-entropy nucleotide region in Figure 2.
